# Guidelines for alternative polyadenylation identification tools using single-cell and spatial transcriptomics data

**DOI:** 10.1101/2024.11.29.626111

**Authors:** Qian Zhao, Magnus Rattray

## Abstract

**Background:** Many popular single-cell and spatial transcriptomics platforms exhibit 3’ bias, making it challenging to resolve all transcripts but potentially more feasible to resolve alternative polyadenylation (APA) events. Despite the development of several tools for identifying APA events in scRNA-seq data, a neutral benchmark is lacking, complicating the choice for biologists.

**Results:** We categorized existing APA analysis tools into three main classes, with the alignment-based class being the largest and we further divided this category into four sub-types. We compared the performance of methods from each algorithmic subtype in terms of site identification, quantification, and differential expression analysis across four single-cell and spatial transcriptomic datasets, using matched nanopore data as ground truth. No single method showed absolute superiority in all comparisons. Therefore, we selected representative methods (Sierra, scAPAtrap, and SCAPE) to deeply analyze the impact of different algorithmic choices on performance. SCAPE which is based on the distance estimation demonstrated less sensitivity to changes in read length and sequencing depth. It identified the most sites and achieved high recall but does not account for the impact of alternative splicing on site identification, leading to a loss in precision. Sierra that fits a coverage distribution is sensitive to changes in sequencing depth and identifies relatively fewer sites, but it considers the impact of junction reads on site identification and this results in relatively high precision. scAPAtrap combines peak calling and soft clipping, both of which are sensitive to sequencing depth. Moreover, soft clipping is particularly sensitive to read length, with increased read length leading to more false positive sites. Quantification consistency was affected by Cell Ranger versions and parameters, influencing downstream analysis but having less effect on differential expression between cell types.

**Conclusions:** Each method has unique strengths. SCAPE is recommended for low-coverage data, scAPAtrap for moderate read lengths including intergenic sites, and Sierra for high-depth data with alternative splicing considerations. Filtering low-confidence sites, choosing appropriate mapping tools, and optimizing window size can improve performance.

## 1 Introduction

Alternative polyadenylation (APA) events are widespread in both animals and plants, directly impacting the stability, localization, and functionality of mRNA by influencing the combination of terminal exons and 3’UTRs. 10x Chromium (single-cell RNA-seq, scRNA-seq) and 10x Visium (spatially resolved RNA-seq, spRNA-seq) from 10x Genomics employ 3’-tagged sequencing which means they are biased towards the 3’ end of transcripts. While this 3’ bias makes it challenging to resolve alternative transcripts for some genes, it is potentially less problematic for resolving APA events.

As these technologies produce reads near the polyA sites rather than directly on them, several computational tools have been developed to infer polyA sites from scRNA-seq data using different rationales. Based on their different purposes and principles, they can be categorized into three main classes (**Figure 1A**, **Table 2**). The first category of methods follows the conventional RNA-seq analysis workflow, based on sequence alignment software such as CellRanger or STARsolo to obtain BAM files. Subsequently, these BAM files are used as input to identify polyA sites or intervals. This category encompasses the largest number of tools and can be further divided into four subcategories. The first subcategory includes scAPA, SCAPTURE, and Sierra, which identify polyA sites by fitting the distribution of reads on the genome or transcriptome (G.-W. Li et al., 2021; Patrick et al., 2020; Shulman and Elkon, 2019). The second subcategory, represented by SCAPE and MAAPER, predicts precise sites by estimating the distance from reads to polyA sites (W. V. Li et al., 2021; Zhou et al., 2022). The third subcategory assumes that the sum of all isoforms from the same gene can explain changes in coverage. Therefore, after identifying the interval of the mapped reads or the furthest polyA site, the remaining sites can be inferred based on change points in the coverage without the need for distribution fitting. Representative methods in this subcategory include scAPAtrap and scDaPars (Gao et al., 2021; Wu et al., 2021). The tools in the fourth subcategory are based on 3’ end soft-clipped reads which potentially contain part of the polyA tail, and the junction of matched sequence and soft-clipped sequence is considered as the inferred polyA site. Representative methods include polyApipe (unpublished, GitHub: https://github.com/MonashBioinformaticsPlatform/polyApipe), tail anchoring from scAPAtrap, and SCINPAS (Moon, Burri, and Zavolan, 2023; Wu et al., 2021). Although polyApipe is unpublished, it can be directly integrated into Seurat using R package PASTA (Kowalski et al., 2023). scAPAtrap consists of two parts: calling peaks based on change points and finding tails based on soft-clipping. The latter can be a filter to correct the sites inferred by the peak calling step, but can also be used independently (Wu et al., 2021).

**Figure 1:**
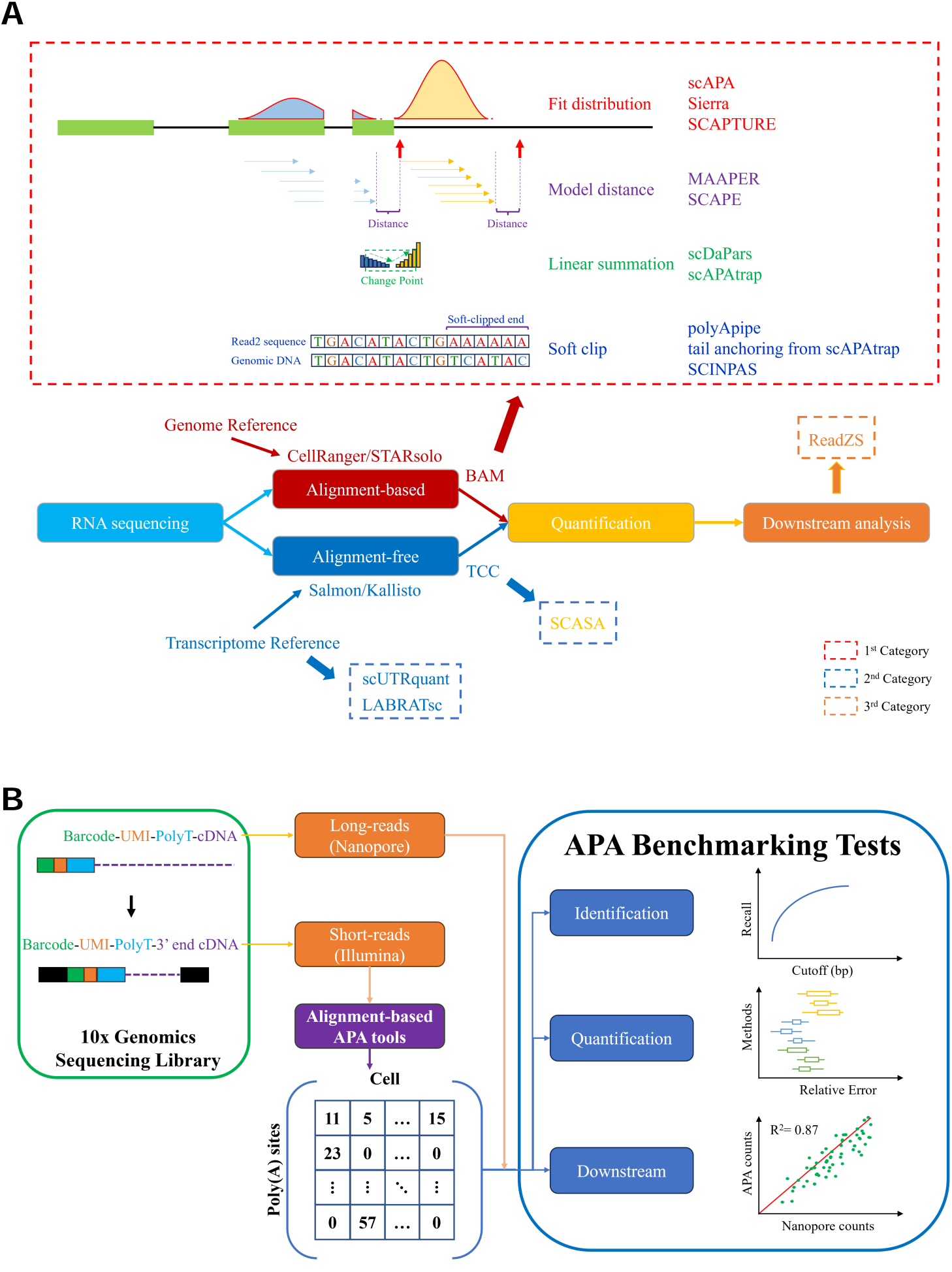
APA identification methods and benchmark workflow. **(A)** Categories of APA identification tools. The first category relies on aligners such as STAR to obtain the distribution of reads on the genome. This category contains the most members and can be divided into four subcategories. The first subcategory identifies polyA signals by estimating whether the coverage satisfies a probability distribution, such as Sierra. The second subcategory infers the position by estimating the distance from read2 to the polyA site, such as SCAPE. The third subcategory infers the sites by finding change points caused by isoform changes, such as scAPAtrap. The fourth subcategory directly identifies reads containing polyA to obtain the position. The second category is based on pseudo-aligners such as Kallisto and can be divided into two subcategories. The first subcategory achieves quantification of isoforms by constructing a special reference. The second subcategory clusters transcripts and quantifies these clusters. The third category does not directly identify sites but focuses on downstream analysis. **(B)** Benchmarking workflow. To compare the performance of different methods, we first ran APA analysis methods and standardized their output into a unified format (sites × cells). Using ONT results as the ground truth, we compared their performance from three perspectives: identification, quantification, and downstream analysis (differential expression analysis).

**Table 1:** Datasets used in this paper.

| Sample name | Type | Cells/Spots | Depth | Read length | GEO |
| --- | --- | --- | --- | --- | --- |
| Mouse Brain E18 (E18) | scRNA-seq | 1159 | 71M | 132 | GSE130708 |
| Mouse Brain E18 (E18) | Nanopore | 1159 | 57M | - | GSE130708 |
| Mouse Olfactory Bulb (MOBV1) | 10X Visium | 918 | 254M | 91 | GSE153859 |
| Mouse Olfactory Bulb (MOBV1) | Nanopore | 918 | 25M | - | GSE153859 |
| Mouse Brain (CBS1) | 10X Visium | 2560 | 217M | 91 | GSE153859 |
| Mouse Brain (CBS1) | Nanopore | 2560 | 38M | - | GSE153859 |
| Mouse Brain (CBS2) | 10X Visium | 2500 | 210M | 91 | GSE153859 |
| Mouse Brain (CBS2) | Nanopore | 2500 | 91M | - | GSE153859 |
| Mouse Olfactory Bulb (MOBC1) | scRNA-seq | 11481 | 547M | 66 | GSE121891 |
| Mouse Brain 1 month (MBM1) | scRNA-seq | 4660 | 327M | 150 | GSE169606 |

**Table 2:** Summary of the performance of each method across various comparisons.

| Method | Subcategory | Identification Precision Rank | Identification Recall Rank | Quantification Mean Rank | DE Mean Rank |
| --- | --- | --- | --- | --- | --- |
| scAPA | 1 | 4 | 6 | 3.5 | 6 |
| Sierra | 1 | 1 | 3 | 3.5 | 4 |
| SCAPTURE | 1 | 2 | 2 | 3.5 | 1 |
| SCAPE | 2 | 6 | 1 | 3 | 1.5 |
| scAPAttrap | 3 | 5 | 5 | 3.5 | 3 |
| polyApipe | 4 | 3 | 3 | 4 | 4 |

The second category leverages pseudo-aligners, such as salmon or kallisto, to perform transcriptome level quantification, indirectly enabling the study of APA events, and includes methods such as scLABRAT, scUTRquant, and SCASA (Fansler, Mitschka, and Mayr, 2023; Gillen, Goering, and Taliaferro, 2021; Goering et al., 2021; Pan et al., 2022). For example, LABRAT constructs a reference for isoform quantification by retaining the 3’ end of transcripts, improving the quantitative effect of the 3’ end, and comparing the differences in APA events by calculating the relative usage ratio (Gillen, Goering, and Taliaferro, 2021; Goering et al., 2021). By constructing the UTRome based on Microwell-Seq, scUTRquant also falls into this category (Fansler, Mitschka, and Mayr, 2023). Unlike the two methods above, SCASA doesn’t optimize the reference. Instead, it focuses on enhancing transcript-level quantification by clustering transcripts to reassign multi-mapping reads (Pan et al., 2022).

A third category of method is represented by ReadZS, which segments the genome into distinct windows, analyzing regions exhibiting differences at the single-cell level to explore differential APA events (Meyer et al., 2022). Once differential regions are identified, Gaussian Mixture Models (GMM) are employed to split peaks and then annotated sites (Meyer et al., 2022). Therefore, unlike other methods, it first identifies differential genomic regions and then identifies sites within those regions.

Conducting benchmark testing for these tools poses a challenge due to their differing principles and design objectives, as well as limitations in available ground truth data. Zhou et al. (2022) developed SCAPE and used simulated and real data to compare SCAPE with other methods in the first category for site identification, demonstrating SCAPE outperforms other methods based on bulk-level Oxford Nanopore Technologies (ONT) data as the ground truth. Due to cellular heterogeneity, bulk-level sequencing misses genes and transcripts that are only expressed in a small subset of cells. Therefore, data from single-cell level ONT may provide better ground truth for benchmarking. Ye et al. (2023) systematically reviewed current methods for predicting or identifying polyA sites using bulk RNA-seq data, scRNA-seq data, and DNA sequence data, and conducted simple benchmark tests on Sierra, SCAPTURE, and scAPAtrap. Their ground truth data was sourced from GENCODE v39, PolyASite 2.0, and PolyA DB 3, which also relied on bulk-level sequencing. The benchmarking primarily emphasized the quantity and distribution of overlapping sites, along with the pattern of polyA signals. Recently, Bryce-Smith et al. (2023) conducted a comprehensive benchmark study for APA identification tools in bulk RNA-seq data. They reviewed 17 APA identification tools and performed benchmark tests on eight of them, comprising both peak-based and pseudo-aligner-based tools. Using paired 3’ end data, they not only conducted analyses for site identification but also carried out tests for quantification. Although this benchmark study encompassed numerous methods, they focused on bulk-level sequencing research, rather than single-cell sequencing. Moreover, while they compared identification and quantification, their study didn’t consider further downstream analyses and they failed to account for the influence of various sequencing metrics on identification capabilities, such as coverage, read length, and junction reads.

It is important for biologists to choose appropriate methods for their research purposes, but there is a lack of independent benchmarking of APA event detection methods at the single-cell level and few applications to spatial transcriptomics data. Here, we aim to provide guidelines to help researchers select the best APA tools based on our benchmarking workflow. We employed representative single-cell level APA identification tools on single-cell and spatial transcriptome sequencing data and validated them using paired ONT data.

## 2 Results

### 2.1 The APA benchmark workflow

Current single-cell APA identification tools vary significantly in their design principles and often differ in usability. These tools can be categorized into three main classes shown in **Figure 1A**. To guide the selection of APA identification tools we have established a benchmarking workflow to evaluate their performance (**Figure 1B**). The workflow is divided into two main parts. The first part involves implementing various APA identification methods and standardizing their outputs into a unified format, namely the polyA sites matrix (polyA sites by cells matrices). The second part compares the results of these tools in terms of identification, quantification, and downstream tasks (differentially expressed APA analysis).

These tools were developed based on 3’ end single-cell sequencing data, so we believe they can also be applied to spatial transcriptomics data generated by 3’ end sequencing, despite the current lack of extensive research in this area. Therefore, apart from single-cell data, we have also implemented and tested these methods on 10x Visium data. The matrix generated from 10x Visium data is polyA sites by spots, rather than sites by cells. Unlike existing benchmark studies that use bulk-level sequencing data as the ground truth, we collected long-read sequencing data from ONT matched with single-cell and spatial transcriptomic short-read sequencing as our ground truth. The shared barcodes and UMIs between Illumina short-read data and ONT data allow us to compare the performance of these tools at a higher resolution. To enhance comparability, only methods capable of generating absolute expression matrices of single-cell-level sites were selected for benchmark testing.

### 2.2 Performance of polyA sites identification among tools

We analyzed the performance of representative APA analysis tools (Sierra, scAPA, SCAPTURE, SCAPE, scAPAtrap, polyApipe) in polyA site identification using four datasets with paired short-read and long-read sequencing data (**Figure 2**, **Table 1**). We transformed the ONT transcript matrix into a polyA site matrix and subsequently compared the predicted site matches with this ground truth (**Figures 2A**–**2B**). The sets of genes identified vary across methods (**Figure S3**). To ensure a fair comparison, we extracted the sites corresponding to genes common to all methods for comparison. Due to the potential distance offset between predicted and true polyA sites, we implemented extended windows on both ends. PolyA sites falling within the overlapping region of the window were considered consistent with the true polyA sites. As the window size increases, the true positive rate grows and then plateaus, resulting in consistent changes in precision and recall (**Figures 2A**–**2B**). For most methods and datasets, the trend slows down and begins to plateau for windows greater than 50nt.

**Figure 2:**
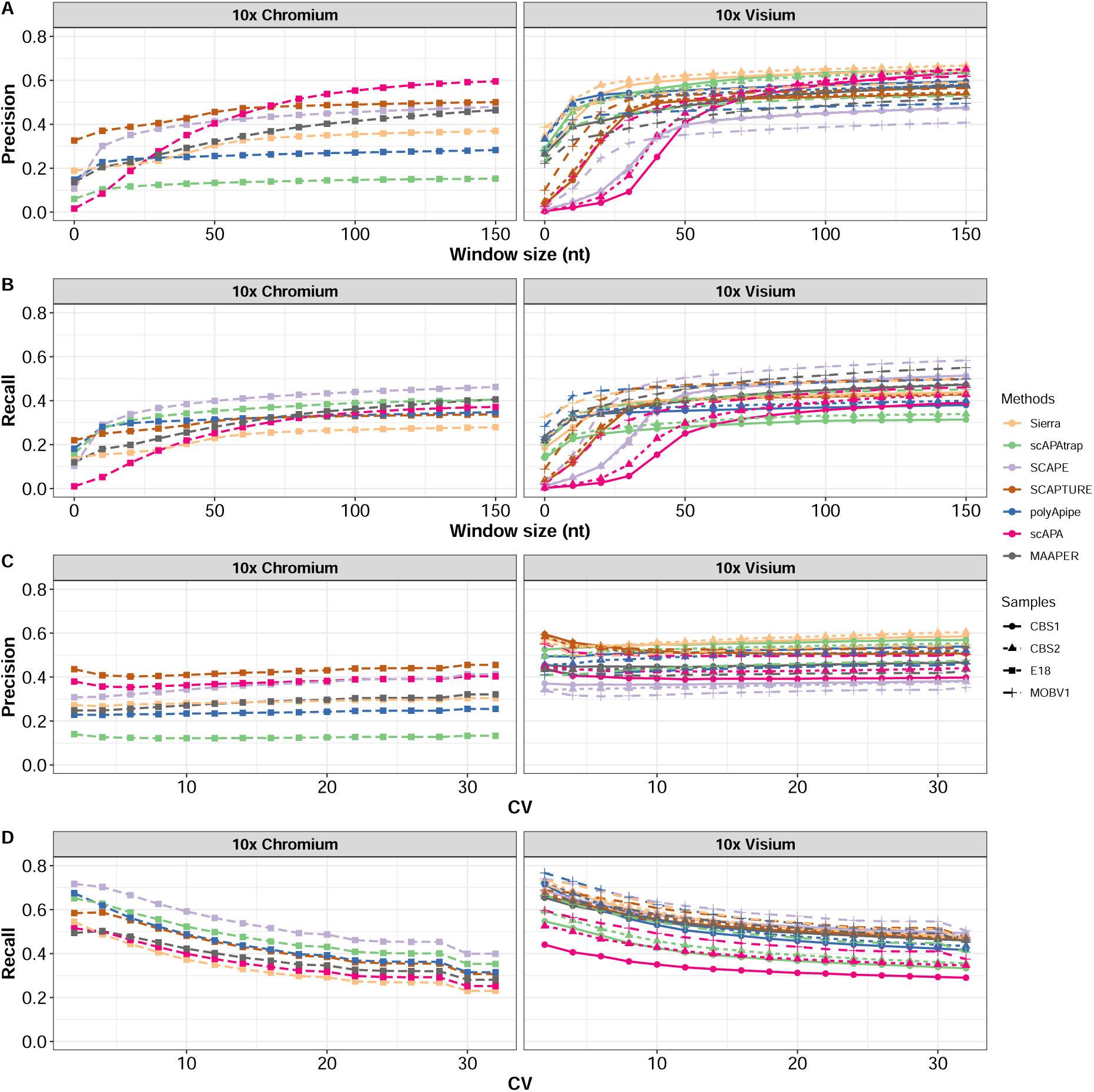
Differences in the performance of polyA site identification. **(A)** Precision of site identification. The x-axis is the window size. As the window increases, the number of overlaps between predicted sites and ground truth sites will increase. The y-axis represents precision, which is the proportion of true sites among predicted sites. **(B)** Recall of site identification. The x-axis is the window size. The y-axis represents recall, which is the proportion of true positive sites in ground truth sites. **(C)** Variation in Precision with changing CV. We fixed the window size to 50nt and filtered the ground truth sites based on the coefficient of variation (CV). The filtered sites are reused to calculate precision. **(D)** Variation in recall with changing CV. The filtered sites are reused to calculate recall.

Notably, as shown in **Figure S2**, there are two different strategies for calculating true positive sites (Bryce-Smith et al., 2023; Zhou et al., 2022). The first strategy considers both predicted sites as true positives when two predicted sites fall within the window range of a single genuine site. The second strategy, however, counts only one true positive in such cases, viewing both predicted sites as corresponding to the same true site. Since site locations are inferred, positional offsets often occur, leading to identification noise in one-to-many or many-to-many cases. This explains the need for setting such strategies and why the performance of these methods is poor when the window is set to zero, especially for identification methods that do not rely on external annotations.

In terms of results, the first strategy’s limitation is that all inferred sites can become true positive sites, resulting in a precision of 1. However, with the second strategy, as there can be multiple sites corresponding to one ground truth, the precision is typically lower. We adopted the second strategy, resulting in fewer true positive sites identified, which is also why the precision and recall values are not as high. This difference in choice can make it difficult to compare performance results across different published studies.

In the E18 single-cell data (10x Chromium), scAPA achieves the best precision as the window size increases, while SCAPE exhibits the highest recall. When the window size is set to 50 nt, SCAPTURE and SCAPE show better precision than scAPA, with SCAPE still achieving the best recall. In spatial transcriptomics data (10x Visium), the performance of each method is similar, particularly in the CBS1 and CBS2 datasets, which are derived from the same tissue. SCAPE identifies the most sites, sacrificing precision for higher recall compared to the other methods, whereas Sierra demonstrates the best precision.

Methods with similar principles tend to exhibit comparable performance. SCAPE and MAAPER, originating from subcategory 2, both show relatively high recall. Similarly, scAPA-trap and polyApipe incorporate a soft-clip mechanism and demonstrate similar patterns, such as lower precision and higher recall in the E18 dataset, but lower recall and higher precision in other datasets. Among the three methods in the first subcategory, which are all based on coverage fitting, there are considerable differences. Sierra identifies peaks in R based on a Gaussian distribution, while SCAPTURE and scAPA use HOMER to identify peaks based on a Poisson distribution, though with some distinctions. SCAPTURE employs transcriptomic annotations, whereas scAPA operates at the genomic level. SCAPTURE performs identification across all cells collectively, while scAPA identifies peaks separately for each cell type. Nevertheless, similar trends can be observed in their changes.

Long-read sequencing has advantages over short-read sequencing in identifying transcript structures, making it well-suited as a ground truth for studying transcript structure variation. However, long-read sequencing also has limitations, such as lower sequencing depth and quality compared to short-read sequencing. Although it provides richer structural and positional information, insufficient depth or quality can decrease the reliability of the identified transcripts, potentially resulting in true positive sites being missed or false positive sites appearing due to alignment errors or low-confidence sites supported by only a few or even a single read. Thus, filtering is necessary to increase the credibility of the ground truth. Given the use of different datasets, we opted for a flexible metric, the coefficient of variation (CV), rather than setting a fixed read threshold. The CV is calculated as the standard deviation of isoform expression across cells divided by the mean expression. This can be applied to all cells or specific cell types. As the CV decreases, low-confidence sites are removed, leading to fewer total ground truth sites, yet precision does not show a significant decline and even slightly improves in some methods, while recall increases notably (**Figures 2C**–**2D**). This suggests that removing low-quality sites has minimal impact on true positive sites and may even help reduce identification noise.

### 2.3 Factors influencing the number of predicted sites

As previously mentioned, methods with similar principles show comparable identification performance, which varies across datasets. To examine how different factors influence the performance of these methods, we selected three representative methods (Sierra, scAPAtrap, and SCAPE) and analyzed the factors affecting the number of identified sites.

#### 2.3.1 Read length effects

When we applied these tools to different datasets (**Table 1**), significant differences in the number of identified polyA sites were observed. For instance, in the E18 dataset, scAPAtrap yielded the highest number of sites (92,598), much more than Sierra (15,252) and SCAPE (28,367). Similarly, in the MBM1 dataset, scAPAtrap identified a large number of sites (165,092), significantly surpassing other methods such as Sierra (58,074) and SCAPE (46,744). However, in the other three spatial transcriptomics datasets, SCAPE identified the most sites, while scAPAtrap’s identified sites noticeably decreased.

We observed substantial differences in read length and coverage among these datasets, potentially explaining the variation in site numbers across different datasets. When reads are shorter, issues such as multi-mapping may arise, whereas increasing read length faces the tradeoff of increased costs and diminishing sequencing quality. Therefore, read length is often a compromise. According to the recommended workflow of 10x Genomics, Read 2 (R2) lengths are typically 91nt for the v3 kit and the Visium kit, while the v2 kit is generally 98nt. However, due to user customization options, data from different sources in the database may have different lengths, often overlooked by researchers, potentially impacting downstream analysis. The first category of APA analysis methods relies on the distribution, sequence, or positional information of R2, and thus is likely to be affected by changes in read length.

To validate this hypothesis, we collected the MBM1 dataset, constructed using the single-cell sequencing v3 kit, with R2 having a length of 150nt. By truncating the reads, we observed that with shorter read lengths, SCAPE and Sierra show a slight decrease in the number of identified sites, while scAPAtrap exhibits a significant decrease, and the number of sites identified by scAPAtrap in this dataset is significantly higher than other methods (**Figure 3A**).

**Figure 3:**
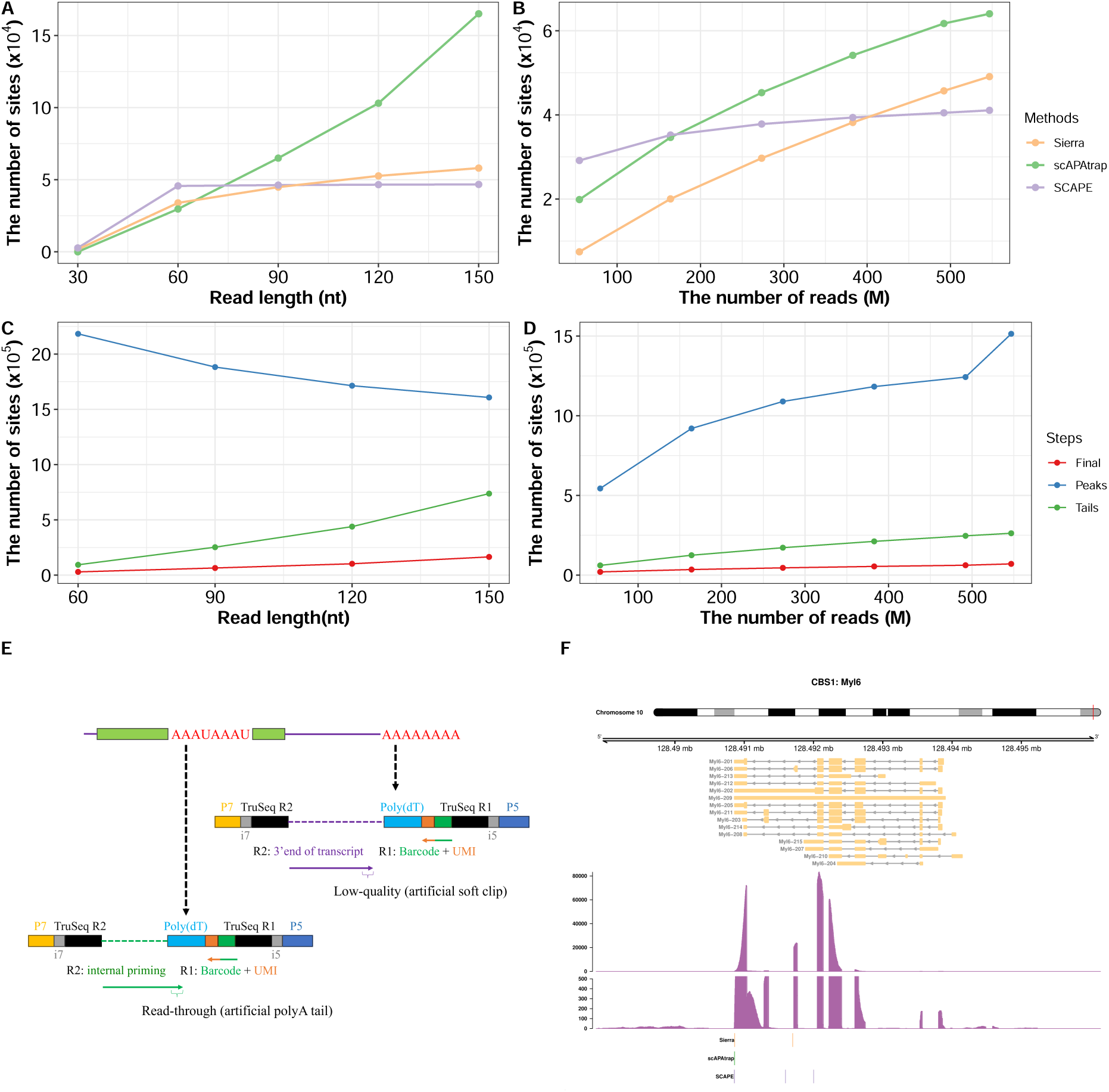
Factors affecting the number of polyA sites. **(A)** Read length truncation analysis. The MBM1 dataset has reads of 150nt length. After reducing the length from 120nt to 30nt, new bam files are generated and the polyA site is re-identified. **(B)** Downsampling analysis. The MOBC1 dataset has more than 500M reads. Different proportions (from 0.9 to 0.1) of reads were randomly sampled. Then, identification was performed on different sampling samples to compare changes in site number. **(C)** The number of sites identified by different parts of scAPAtrap varies with read length. scAPAtrap includes two distinct site identification mechanisms: the first is based on quantile search for change points (blue), and the second is through soft-clipping to search for reads containing polyA tails (green). The intersection of these two methods forms the final result (red). **(D)** The number of sites identified by different parts of scAPAtrap varies with sequencing depth. **(E)** Challenges in identifying sites using soft clip methods. Oligo-dT primer can capture either the polyA tail or A-rich regions within the transcript (indicated in red). The captured transcript fragments are used to construct the library. If the sequencing quality decreases as the Read 2 length increases, it may be mistakenly identified as a soft clip. Internal priming occurs when an A-rich region within the transcript is captured, resulting in shorter fragments. As read length increases, read-through can occur, causing the end of the reads to correspond to the original oligo-dT part and be misidentified as a polyA tail. Both scenarios can lead to an increase in false positive sites. **(F)** Coverage of the Myl6 gene in the CBS1 dataset. At the top is the corresponding chromosomal location, followed by transcripts from the reference annotation, the read coverage (including a subplot ranging from 0 to 500 to zoom in on the low-coverage region), and the locations of the sites identified by the three APA analysis methods.

We further hypothesized that the reason scAPAtrap exhibits more pronounced changes is because it relies on R2 containing the polyA tail for detecting sites, and increasing the length of R2 results in more reads covering the polyA tail. To test this hypothesis, we compared the number of identified sites in the two steps of site identification in scAPAtrap, namely peak find and tail anchor, with increasing read lengths. As shown in **Figure 3C**, as the read length increases, the number of sites identified by tail anchor increases, while the number identified by peak find decreases. Therefore, the significant increase in scAPAtrap sites is due to the increase in sites at the tail anchor step.

The rationale of tail anchor site identification is a component the fourth subcategory of the first category of APA tool (**Figure 1A**, **Table 2**). This approach uses polyA tail sequence information from R2 and is a biologically reasonable strategy, but it has some limitations. First, the premise of this method is the presence of a sufficient number of reads containing part of the polyA tails. Therefore, if the reads are too short or too few, this method may not be applicable, resulting in the loss of signals from many true sites. Second, although increasing the read length results in more polyA fragments being sequenced, it also increases the occurrence of read-through events, especially when internal priming is present. Additionally, the increase in length implies that the sequencing quality at the ends of reads may not be sufficient. Both of these situations can lead to a higher proportion of false positives (**Figure 3E**). Similarly, scAPAtrap identified an unusually high number of sites in the E18 dataset with only 71M reads, which can also be explained by its longer read length of up to 132nt. However, this increased number of sites may predominantly consist of false positives (**Figure 2C**). Besides, the decrease in sites identified by scAPAtrap’s peak finding step as the length increases may be mainly due to its filtering out peaks shorter than the read length. It is important to exercise caution when dealing with reads longer than the standard length, especially when employing soft-clipping mechanisms.

#### 2.3.2 Sequencing depth effects

Coverage is an important factor in RNA-seq analysis. Alignment-based methods generally require a sufficient number of reads as a prerequisite, although different methods may be affected to varying degrees. We selected the MOBC1 dataset with 547 million reads and conducted a downsampling analysis to validate how changes in coverage affect the identification of sites. When the coverage dropped below 400 million, Sierra obtained fewer sites than SCAPE, and when the coverage dropped below 200 million, scAPAtrap obtained fewer sites than SCAPE (**Figure 3B**). Although the number of sites identified by all three methods decreased gradually as the coverage decreased from 547 million, SCAPE showed a smaller decrease compared to the other methods, demonstrating its more robust performance regarding coverage, consistent with its underlying principle.

We also examined the number of sites obtained by each of the two steps in scAPAtrap (**Figure 3D**). The results showed that sites decreased significantly in both steps when the number of reads was reduced. This is consistent with the rationale that a decrease in the number of reads leads to the disappearance of some low-abundance peaks and makes it more challenging to identify change points between peaks. It also randomly reduces the number of reads containing polyA tails.

#### 2.3.3 Splicing effects

Splicing events occurring at the 3’ end of transcripts can also affect the identification of polyA sites. The presence of introns creates gaps in the genomic coverage distribution. Due to the randomness in library construction, reads from the same transcript may be allocated to the exon before and the exon after an intron. Additionally, some junction reads may be mapped incorrectly, increasing instances of soft clipping. These factors can lead to an increase in false positive APA events.

Among the currently published methods, Sierra uses RegTools to extract junctions, splitting the fitting process into two steps: within-junction and across-junction, thus reducing the impact of splicing (Patrick et al., 2020). SCAPTURE uses HOMER to identify peaks on the transcriptome rather than the genome, thereby avoiding this issue (G.-W. Li et al., 2021). However, because it relies heavily on transcriptome information, it may have a weaker capability in identifying novel sites, such as disease-associated intronic polyadenylation sites (IPA). False positive sites induced by the splicing process are often overlooked and lack systematic evaluation methods.

For biologists, the focus may often be on one or a few genes from the same family or signalling pathway, so even if the number of splice-affected sites is small, they may still be crucial in specific studies. To detect the effects of splicing and junctions, we selected representative genes, such as Myl6, and observed whether sites were influenced by the presence of junctions through coverage plots. As shown in **Figure 3F** and **Figure S6**, all three methods were affected to varying degrees when identifying sites across the Myl6 gene body. This suggests that when an intron is detected in the 3’ end of a gene, extra attention should be paid to whether the identified polyA sites are affected by splicing.

#### 2.3.4 Differences in number and location of polyA sites across methods

Next, we compared the distribution of sites generated by different methods through annotation of the predicted sites (**Figures S4**–**S5**). From **Figure S4** it can be observed that the number of single-site genes is the highest. For the MOBV1, CBS1, and CBS2 datasets, the median number of genes predicted by Sierra and scAPAtrap is 1, indicating that more than half of the genes are predicted to have only one site by these methods. However, for SCAPE in these three datasets, the median is 2, identifying more multi-site genes. In the E18 dataset, Sierra has a median of 1, SCAPE has a median of 2, and scAPAtrap has a median of 3. It can also be seen from the figure that the number of genes with more than 6 sites in scAPAtrap is only second to the number of single-site genes, which may be attributed to the performance of tail anchor on longer read lengths as mentioned earlier.

According to the position of the sites on the gene, they can be divided into 3’ UTR, Exon, Intron, regions within 1 kb downstream of the last exon (LastExon1Kb), regions within 1 kb downstream of the 3’ UTR (3UTR 1kb), regions within 2 kb downstream of the 3’ UTR (3UTR 2kb), and intergenic regions (INTERGENIC) (**Figure S5**). Sierra’s identification range mainly focuses on the transcriptome and junction corresponding regions, so the sites are primarily concentrated in UTRs, exons, and introns. In all four datasets, there is a phenomenon where the number of sites from the 3’ UTR decreases as the number of sites increases, while the number of sites from introns increases. The performance of scAPAtrap is similar to Sierra, except that it includes a large number of sites located in intergenic regions, consistent with its whole-genome-wide site identification function. Due to the lack of corresponding gene annotation, the number of sites contained in the INTERGENIC category exceeds 6 and is assigned as “6+”. Since the number of sites identified in the three spatial transcriptomics datasets is relatively small, the proportion of sites from intergenic regions will be more significant. SCAPE shows a more uniform distribution across all datasets, mainly concentrated in the 3’ UTRs, with low proportions of other features, such as introns, which may be due to its built-in mechanism for removing sites in internal priming regions.

Internal priming refers to the process in which the A-rich region in RNA or cDNA is mistakenly recognized as polyA by oligo-dT primer for amplification. This process results in a certain amount of truncated cDNA in the sequencing library, which is considered the primary source of intronic reads. Reads generated from internal priming interfere with the identification of polyA sites and can increase false-positive sites, especially false IPA. Although a high proportion of intron sites may indeed indicate interference from internal priming, it is important to note that the analysis of genomic features relies heavily on current annotations, and annotations are usually assigned based on user-defined priorities, which may not reflect the actual situation. For example, there may be an isoform whose 5’ UTR region coincides with the intron or exon region of another isoform, or an isoform’s intron or exon region coincides with the 3’ UTR region of another isoform, all of which can affect the composition ratio.

The conventional method for eliminating such sites or reads involves detecting the presence of an A-rich region downstream. Sierra provides annotation functions to search for the presence of A-rich regions, allowing users to decide whether to include these sites. SCAPE employs built-in functions to preemptively remove certain intronic sites. Excessive removal of intronic sites or those near A-rich regions may result in the loss of biologically significant information.

### 2.4 Performance of polyA sites quantification among tools

For alignment-based methods, it is usually necessary to identify sites from the BAM file and then quantify the identified sites or regions for downstream analyses. Downstream analyses such as clustering, differential expression, and differential usage rely heavily on quantification. Therefore, we believe it is necessary to consider the potential differences in quantification among different tools when selecting an appropriate APA analysis tool.

We calculated the correlation coefficients between matched predicted sites and ground truth sites in the overlapping gene set among all methods. As shown in **Figure 4A**, as the window size increases, the number of overlapping sites increases, the correlation coefficient begins to fluctuate, and the relative error tends to decrease. Once the number of sites reaches a plateau, the influence of further increases in window size diminishes.

**Figure 4:**
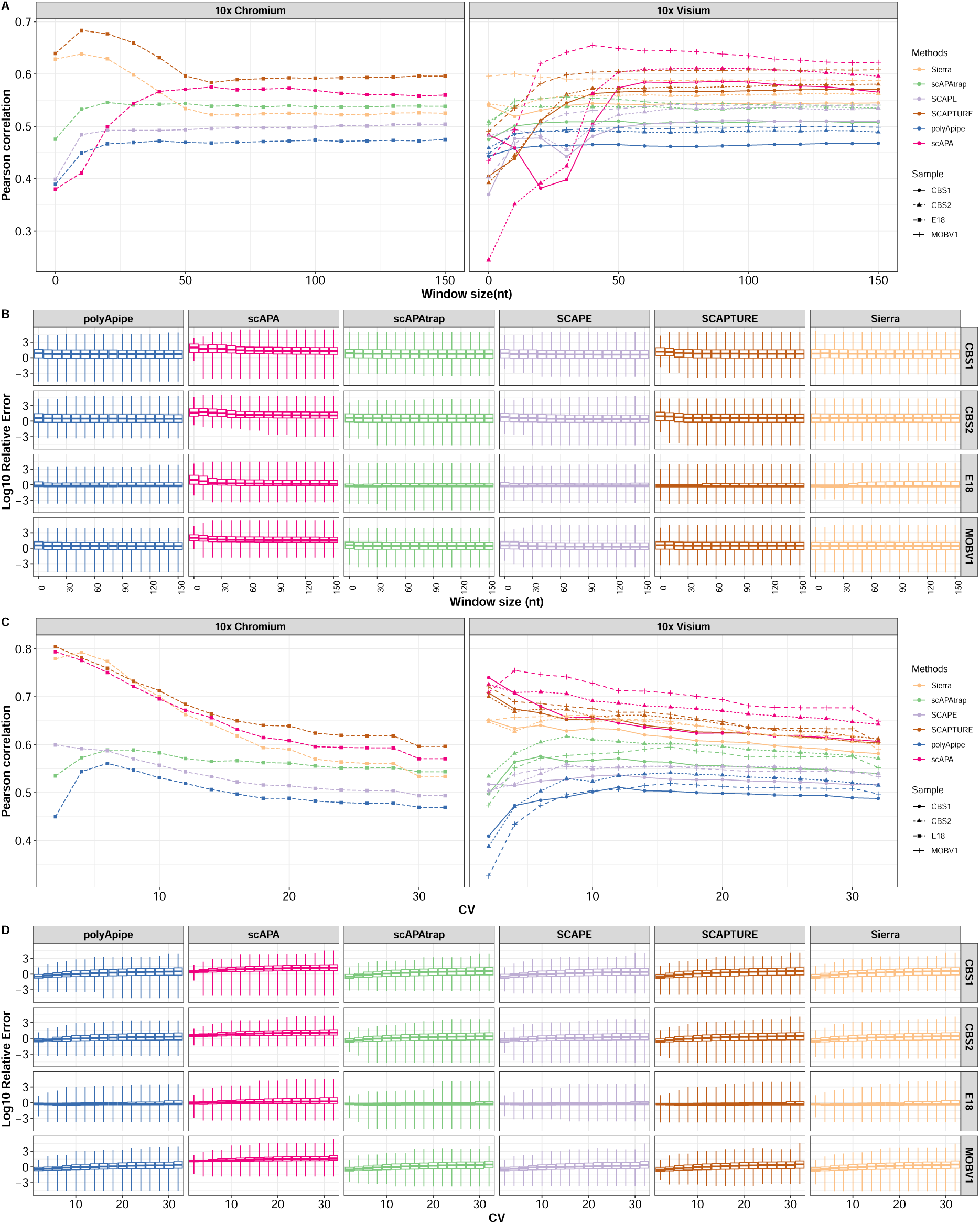
The performance of quantification. **(A)** Site-level correlation. The x-axis represents window size, and the y-axis represents the Pearson correlation coefficient. The log-normalized expression values of predicted sites and their matched ground truth sites are used to calculate the correlation coefficient. Only the site with the highest expression is retained among multiple sites falling within the same window. The window size ranges from 0 to 150, with intervals of 10. **(B)** Site-level RE. The x-axis represents window size, and the y-axis represents the log-transformed relative error (RE). **(C)** Variation in correlation with changing CV. We fixed the window size to 50nt and filtered the ground truth sites based on the coefficient of variation (CV). The filtered sites are reused to calculate Pearson correlation. **(D)** Variation in RE with changing CV. The filtered sites are reused to calculate RE.

Increasing the window size allows more predicted sites to match true sites, leading to an increased number of identified sites, typically accompanied by a generally rising correlation coefficient, though there are exceptions. For instance, in the single-cell E18 dataset, Sierra and SCAPTURE show a declining trend before reaching a plateau, whereas scAPA, belonging to the same subclass, shows an upward trend. When the window size is zero, scAPA identifies the fewest true positive sites due to coordinate shifts, resulting in the lowest precision and recall. Additionally, coverage-based methods generally require a minimum coverage threshold, since scAPA analyzes each cell type’s BAM file individually, low-expression sites are often difficult to detect, especially using low-depth dataset E18. Consequently, the scatter plot reveals an absence of low-expression APA sites, with only a limited number of medium-to-high expression sites, resulting in an inverted-J shape distribution and a very low correlation coefficient (**Figure S9**). As the window size increases, more sites are included, particularly those with medium-to-high expression, expanding the dynamic range and significantly increasing the correlation coefficient (**Figure S10**). Sierra and SCAPTURE face the similar coverage threshold issue, also displaying an inverted-J shape in the scatter plots. Unlike scAPA, however, they identify more sites when the window size is zero, particularly including those with medium-to-high expression levels, yielding a higher correlation coefficient. As the window size increases, more sites with high expression in APA analysis but medium-to-low expression in ONT data are included, leading to a decline in the correlation coefficient.

Besides, incorrect assignments may cause a decrease in correlation coefficients. Incorrect assignments occur due to the current lack of a mechanism, which allows multiple nearby predicted sites to be accurately matched to ground truth sites with consistent expression levels. To avoid additional bias, when multiple sites are matched, we retain only the site with the highest expression level, as this site is most likely to be identified. However, this can also lead to low-expressing predicted sites being assigned to high-expressing ground truth sites, causing a decrease in correlation.

When we used another metric, relative errors (RE), we observed that within the 0 to 50nt window range, the median RE shows a decrease (**Figure 4B**). Due to the imbalance in the first subcategory, sites with high APA counts and low ONT counts increase the RE. Thus, even when these three methods show high correlation coefficients, it is still possible to exhibit relatively poor median RE, especially scAPA.

Despite the generally lower sequencing depth of ONT compared to Illumina, and its higher sequencing error rate making it less suitable for quantitative analysis, we still selected ONT data for quantitative benchmarking. This decision was based on the fact that the ONT data we collected shared barcodes and Unique Molecular Identifiers (UMIs) with the matched Illumina data from the same library, ensuring high consistency (**Figure S7**). Additionally, the advantage of long read lengths in transcript identification allows for better transcript quantification, reducing the probability of false positives. To circumvent issues related to ONT sequencing depth, we used CV to remove polyA sites with low signal-to-noise, similar to identification. We set a CV threshold to retain high-confidence sites and then re-evaluated the quantification performance of the APA analysis methods (**Figures 4C**–**4D**). The underlying assumption posits that low-confidence nanopore sites may not represent genuine sites with consistent count numbers. This inconsistency could lead to discrepancies in predicted sites, resulting in anomalous correlation coefficients.

As the CV decreases, high-confidence sites are retained, and less noise may lead to more reasonable correlation coefficients, potentially resulting in increased correlation. An improvement in correlation was observed in all methods, with methods from the subcategory I showing the best performance. However, there are still some methods that show a decrease in correlation coefficient as CV decreases. For example, there was a gradual decrease after a slight increase in correlation with decreasing CV in all spatial transcriptomics datasets for SCAPE. There are three possible reasons for this phenomenon. First, as the CV decreases, a reduced dynamic range of sites leads to a decrease in the correlation coefficient. Second, SCAPE identified the highest number of sites among the three spatial transcriptomics datasets. Low-quality sites derived from ONT might have incidentally matched the sites identified by SCAPE. When these sites were filtered out, the correlation decreased. Third, the positions of the sites inferred by SCAPE may deviate from the true positions, and the lack of a site alignment mechanism might exacerbate mismatches when a large number of sites are removed from the long-read data. Similarly, in all methods, the relative error decreases as the CV is reduced (**Figure 4D**).

Although we used the overlapping gene set to ensure fairness and facilitate parallelization in the pipeline for improved efficiency, we further aimed to achieve fairer comparison results by reducing the impact of site identification differences. This resulted in the same set of polyA sites for calculating correlation coefficients and RE. As shown in the **Figure S11**, these methods display a similar increase-to-plateau trend, with higher maximum correlation coefficients and smaller differences between methods. While scAPA performs best in CBS1 and CBS2, it underperforms in E18 and MOBV1, whereas SCAPE demonstrates consistently strong and stable performance across all datasets. In terms of RE, scAPA still performs poorly, whereas the differences among other methods are minimal. This indicates that scAPA still has shortcomings in quantification through the generated annotations (**Figure S11**).

### 2.5 Factors influencing quantification steps

#### 2.5.1 Site quantification tools

We further analyze the factors contributing to the differences in quantification performance. We found scAPA showed a high correlation coefficient but a very poor relative error (RE), and it consistently performed worse than other methods across the three tests (**Figure 4D** and **Figure S11**). Although scatter plots revealed biases in the first subcategory, this was insufficient to explain why scAPA had abnormally high relative error.

To further investigate the cause, we calculated the correlation coefficient using the same set of polyA sites and examined their distribution through scatter plots (**Figure S11** and **Figure 5A**). The results showed that even with the same ONT sites, the distribution of scAPA shifted significantly to the right. This indicates that while the ONT expression levels were consistent, the counts quantified by scAPA were significantly higher than those of other methods. The likely reason for this issue is the difference in quantification methods and tools. Among the six methods compared, Sierra and SCAPE used R and Python code, respectively, to identify non-redundant UMIs, while the remaining four methods used featureCounts (**Figure 5B**). The key difference is that, except for scAPA, the other methods used a combination of featureCounts and UMI-tools. This suggests that scAPA might have quantified reads instead of UMIs, leading to an overall bias in scAPA’s quantification results.

**Figure 5:**
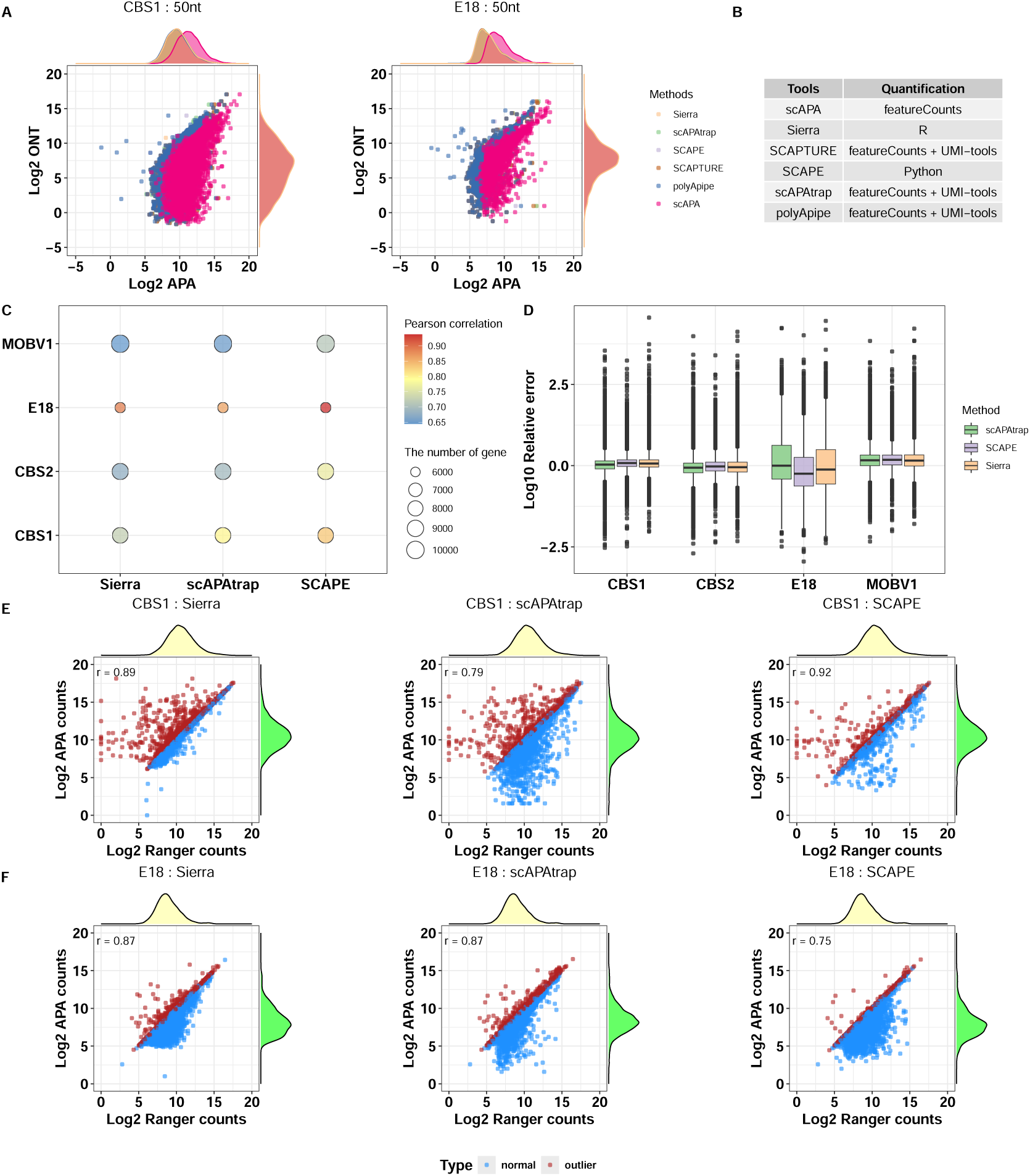
Factors affecting the quantification of polyA sites. **(A)** Scatter plot of the same set of sites among different methods. **(B)** Summary of quantification tools. **(C)** Gene-level correlation. Identified sites were annotated, and the counts from sites originating from the same gene were summed to obtain gene expression levels. Overlapping genes were then selected for comparison with ONT expression levels. The size of the bubbles represents the number of overlapping genes, while the colour intensity indicates the strength of the Pearson correlation coefficient **(D)** Gene-level RE. The relative error (RE) between gene expression levels identified by the APA analysis tools and ONT gene expression levels. The x-axis represents the samples, and the y-axis represents the logarithm of the RE. **(E)** Correlation between gene expression levels obtained by APA analysis methods and those obtained by Space Ranger. Each point represents a gene, with the x-axis showing the log-transformed gene expression levels from Space Ranger, and the y-axis showing the log-transformed expression levels obtained by the APA analysis tool. Red dots indicate outliers, where the APA tool’s expression levels exceed those of Space Ranger, while blue dots represent normal data. **(F)** Correlation between gene expression levels obtained by APA analysis methods and those obtained by Cell Ranger. Each point represents a gene, with the x-axis showing the log-transformed gene expression levels from Cell Ranger, and the y-axis showing the log-transformed expression levels obtained by the APA analysis tool. Red dots indicate outliers, where the APA tool’s expression levels exceed those of Cell Ranger, while blue dots represent normal data.

#### 2.5.2 Gene level quantification

The accuracy of site quantification depends on the accuracy of gene quantification. If the gene expression level obtained by the APA analysis is significantly lower than that of Cell Ranger, for example, due to the loss of a large number of reads, this would lead to poor site quantification results. Therefore, we explored whether the gene-level expression generated by APA analysis tools correlates well with the quantification results from ONT. In **Figure 5C** and **Figure S7**, the number of genes identified in the E18 dataset is the smallest, with correlation values generally higher than those observed in the other three datasets. Among the three spatial transcriptomics datasets, CBS1 exhibits the fewest genes and the highest overall correlation, whereas MOBV1 identifies the most genes and demonstrates the lowest correlation. SCAPE consistently shows the strongest correlation with ONT across all datasets, while Sierra and scAPAtrap perform comparably. Regarding relative error (RE), the three spatial transcriptomics datasets exhibit comparable performance across the three methods (**Figure 5D**). SCAPE shows a slightly larger median than the other two methods, making it the weakest performer overall. However, in the E18 dataset, SCAPE outperforms the others, while scAPAtrap performs the worst.

The performance of gene quantification is influenced by both the method used and the number of genes included in the dataset. In the E18 dataset, although all methods exhibited high correlation, this dataset is characterized by a relatively shallow sequencing depth and longer read lengths. As a result, the scAPAtrap method identified significantly more sites compared to the other two methods. Consequently, while scAPAtrap did not demonstrate the best performance at the gene level, it gained a certain advantage when calculating the correlation of overlapping sites.

#### 2.5.3 Alignment method

Besides the factors affecting site identification and gene quantification, the alignment step also influences site quantification. APA analysis tools re-quantify the sites based on the generated annotation files for those sites. This means that some reads may not be assigned to any site, although they can map to gene annotations. There may also be other issues, such as duplicated assignments. Therefore, it is worth verifying whether the gene expression levels identified by APA analysis tools are consistent with the quantification results from Cell Ranger or Space Ranger. Theoretically, using BAM files generated by Cell Ranger or Space Ranger as input for site identification should not result in gene expression levels, obtained by summing up sites from the same gene, that exceed those obtained by Cell Ranger or Space Ranger.

We compared the UMI counts of genes quantified by Cell Ranger or Space Ranger with the identified sites by APA analysis tools (**Figures 5E**–**5F** and **Figure S8**). At the cell level, they showed high similarity. However, it can be observed that in the E18 data using the new version of Cell Ranger (version 7.0.1), there are very few outliers (defined as counts obtained by APA analysis tools exceeding those of Cell Ranger or Space Ranger). In contrast, outliers predominate in the MOBV1 data using the new version of Space Ranger (version 2.0.1). When comparing at the gene level, although *R*^2^ looks good, many outliers remain (**Figures 5E**–**5F**). Similarly, the number of outliers in the spatial transcriptomics data is significantly higher than in the single-cell data.

The occurrence of these outliers is related to the version of Cell Ranger. Starting from version 6.0, Cell Ranger uses the intron mode by default during quantification, a parameter that did not exist in the Space Ranger. This is why there are significantly more outliers in 10x Visium data than in single-cell data. Additionally, starting from version 3.0, Cell Ranger introduced an extra filtering tag “xf” in the BAM files, which is often not accounted for by APA analysis tools. This is another reason why there are still outlier sites in single-cell data. It is worth noting that if the APA analysis methods developed for single-cell or spatial transcriptomic data do not account for the software version, the newly identified clusters or differentially expressed APA isoforms in downstream analysis may be due to the lack of intronic reads or the use of different filtering tags, rather than providing information beyond gene expression levels.

### 2.6 Perfomance of analyzing differential polyA sites

Quantification of gene expression or polyA site usage can be affected by technology-specific biases that could change the relative expression of genes or APA sites within a sample. However, biologists are more often interested in changes of the same gene or site across biological samples, cell types or spatial regions. In this case, it should be more straightforward to compare results between different technologies, since we expect biases to have less impact when comparing expression of the same gene or APA site.

We conducted differential expression analysis at the site level. We selected the two cell types with the highest cell counts in each dataset and used the DESeq2 method to identify differentially expressed sites between these two cell populations (FDR *<* 0.05). The differentially expressed sites identified by ONT between the two cell types are consistent across all methods, but coordinate shifts are still present. Thus, as in the previous sessions, different window sizes are applied to these differentially expressed sites to obtain true positives. Since the number of differentially expressed sites detected by ONT is significantly lower than that identified by APA analysis tools, recall rather than precision better reflects the performance of these tools in detecting differential expression. The **Figure 6A** shows a trend where recall initially increases with window size and then gradually reaches a plateau, with the transition point for most methods still being at 50 nt. Overall, the recall of these methods is close. Among them, scAPA and scAPAtrap perform the worst, while SCAPE and SCAPTURE show better and more stable performance. Sierra performs best and most consistently in spatial transcriptomics data but shows weaker performance in single-cell data.

**Figure 6:**
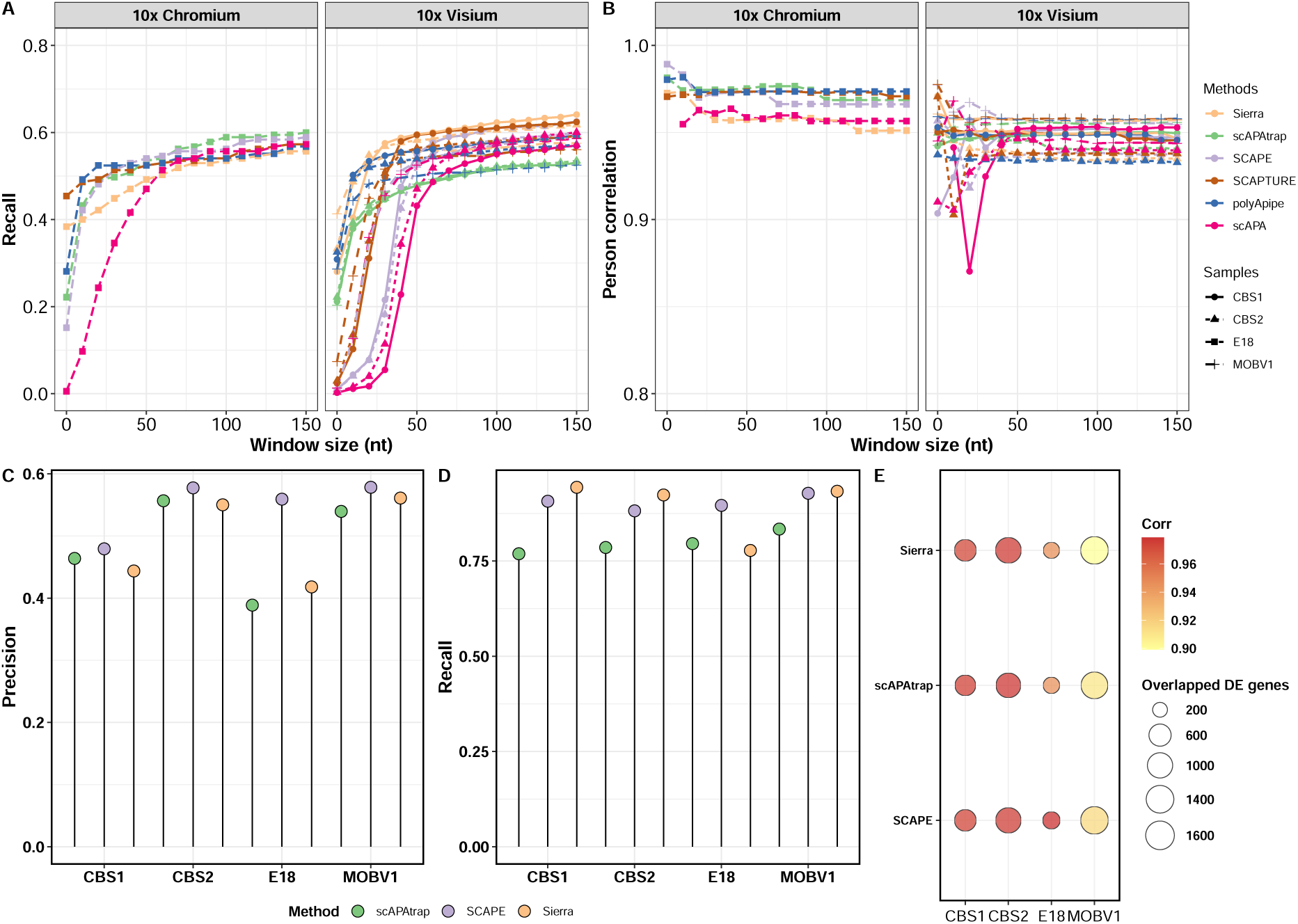
The performance of differential expression. **(A)** Recall of differentially expressed sites. Differentially expressed sites were identified using DESeq2. Overlapping sites with consistent regulation direction were defined as true positive sites and used to calculate recall. The x-axis represents the window size. **(B)** LogFC correlation of differentially expressed sites. The Pearson correlation coefficients between the logFCs of overlapping differentially expressed sites. The colour represents different methods. The x-axis represents the window size, and the y-axis represents the correlation coefficient. **(C)** Precision of differentially expressed genes. Differentially expressed genes are obtained using DESeq2. Genes with consistent regulation direction are considered true positive sites and are used to calculate precision and recall. **(D)** Recall of differentially expressed genes. **(E)** LogFC correlation of differentially expressed genes. The Pearson correlation coefficients between the logFCs of overlapping differentially expressed genes. The colour represents the strength of the correlation, and the size represents the number of overlapping sites. The x-axis represents the four datasets, and the y-axis represents the three methods.

We also compared the correlation of log fold change (LogFC), and although the number of overlapping differentially expressed genes varied across datasets, most correlations were higher than 0.90 (**Figure 6B**). This is consistent with our hypothesis that the results of differential expression analysis may be less influenced by site quantification. This further illustrates that the correlation between the process of differential expression analysis and identification and quantification may not be as evident, suggesting that different downstream analyses may be subject to varying influences.

Similarly, we selected representative methods to examine whether the performance in differential gene analysis at the gene level can explain the performance observed in site-level analysis. The **Figures 6C**–**6E** shows that in terms of precision, SCAPE performed best across all datasets. In CBS1 and CBS2, scAPAtrap slightly outperformed Sierra, but in E18 and MOBV1, Sierra performed slightly better than scAPAtrap. Across all three spatial transcriptomic datasets, scAPAtrap performed the poorest in the recall. In CBS1 and CBS2, Sierra outperformed SCAPE, while in MOBV1, SCAPE and Sierra were comparable. In E18, SCAPE performed the best, followed by scAPAtrap, and then Sierra. The correlations of LogFC were all higher than 0.90. It can be seen that differential expression analysis at the site level is consistent with that at the gene level.

## 3 Discussion

The 3’ bias of read coverage in both 10x Chromium and 10x Visium datasets limits the ability to identify transcripts but may offer advantages for studying APA events. While several methods have been designed for identifying polyA sites in single-cell transcriptomics, few have been applied to spatial transcriptomics, and a neutral benchmark is lacking. We classified existing methods and selected representative tools from each category for benchmarking on both singlecell and spatial transcriptomic data. **Table 2** provides a summary of their performance in site identification, quantification, and differential expression analysis, using paired nanopore data as the ground truth.

We found that methods inferring sites by estimating the distance between reads and polyA, such as SCAPE, exhibit greater robustness to changes in coverage and read length, while soft clip-based methods are exceptionally sensitive to changes in read length and may introduce more false positive sites due to sequencing quality and read-through events. The method scAPAtrap, which combines change point and soft clip approaches to identify sites across the genome, inherits disadvantages from both methods, being sensitive to coverage and read length. Additionally, due to its strategy of taking the intersection of the two methods, scAPAtrap may result in the loss of true positive sites. There is an option with scAPAtrap to use a union strategy, but that leads to a sharp increase in the number of sites and a resulting decrease in precision. Methods like Sierra fit coverage to predict site positions, exhibiting relative insensitivity to changes in read length but sensitivity to changes in coverage.

In terms of site identification, SCAPE shows the best recall. However, the lack of a mechanism for splice awareness and the presence of a mechanism for removing a subset of intronic sites may impact its practical applicability to biological questions, such as analyzing the abundance of IPAs in certain tumour cells. Additionally, since SCAPE is designed for paired-end sequencing methods, its application to more prevalent single-end sequencing may result in more potential offsets in site identification due to the use of empirical parameters, necessitating the selection of an appropriate window size to address this issue. Sierra demonstrates the highest precision on the 10x Visium data rather than the lower depth E18 scRNA-Seq data, highlighting its limitation with respect to sequencing depth.

In terms of quantification, we observed significant discrepancies in the quantitative results obtained by APA analysis tools compared to the gene quantification results provided by Cell Ranger or Space Ranger, depending on the version of Cell Ranger used. This suggests that the identification of novel clusters or markers in downstream analysis may be influenced by whether Cell Ranger employs new filters and includes intronic reads, rather than solely by the additional information provided by APA events themselves. The comparison of site quantification utilized two metrics: the correlation coefficient and relative error. Methods based on similar principles exhibited similar trends; however, some methods, particularly scAPA, showed high correlation but very poor relative error. This discrepancy is caused by two factors. The first reason is that methods in the first subclass require a certain number of reads to fit the coverage, leading to bias in site identification. The second reason is the inappropriate choice of quantification tools, such as using featureCounts to quantify read numbers without combining it with UMI-tools to quantify unique UMI counts.

Differential expression analysis considers the fold change of genes under different conditions, which may be relatively unaffected by absolute quantification. Therefore, we chose differential expression analysis as a representative downstream analysis. We found that in gene-level differential expression analysis, SCAPE performed well in precision, while Sierra excelled in recall. Additionally, the correlation of LogFC is high for both methods. In our analysis of sites, we also observed a transition point at 50 nt. Considering the entire window range, SCAPE performed robust across all datasets in terms of correlation and recall. However, if only the 50 nt position is considered, Sierra outperformed SCAPE in the three spatial transcriptomic datasets. These results are consistent with our hypothesis that different downstream analyses are influenced differently by the performance of identification and quantification.

Although we compared the performance of identification, quantification, and differential analysis, the current results do not support any single method dominating all datasets and tests. Therefore, when using these methods, it is essential to consider one’s experimental objectives. For instance, for datasets with low sequencing depth, methods like SCAPE may be more suitable. Meanwhile, for situations necessitating consideration of alternative splicing or differential transcript usage, particularly with coverage exceeding 200M, Sierra may be more appropriate. If the sequencing read length exceeds the standard length, then soft-clip-based methods may not be applicable.

Developing a benchmarking approach for poly-A site detection methods is challenging for several reasons. We have attempted to minimise biases in our metrics as much as possible but challenges remain. We selected long read benchmark data that was paired with the analysed short read data at the cell or spot level through shared barcodes, enabling a higher resolution ground truth and avoiding issues of cellular heterogeneity. However, the effectiveness of singlecell nanopore analysis tools and metrics such as sequencing read length and saturation still affect the validity of our ground truth. Furthermore, limitations in the sequencing depth of the nanopore data meant that we had to filter out poly-A sites from the ground truth to ensure reliable sites, inevitably leading to a loss of genuine sites and an underestimate in the precision metric due to some true positives being reported as false positive. Therefore, it remains worthwhile to explore a more effective integration of short-read and long-read quantification tools, as well as a more reasonable ground truth. Similarly constrained by sequencing depth, current computational methods typically use all reads as input for prediction, potentially missing some cell-type-specific signals. Additionally, the lack of a systematic evaluation of sites affected by junctions and internal priming is another important issue, and the identification and removal of internal priming signals is a worthwhile direction for future methodological developments. In downstream analysis, clustering, enrichment analysis, and differential UTR usage also await further study.

## 4 Conclusions

We conducted a benchmark analysis of various APA analysis tools in terms of site identification, quantification, and differential expression analysis. No single method outperformed others across all comparison metrics. Methods based on similar principles exhibited similar performance. The first subclass, which relies on coverage fitting, the third subclass, based on change points, and the fourth subclass, based on soft-clipping, were more sensitive to sequencing depth compared to the second subclass, which relies on distance estimation. Additionally, methods in the fourth subclass were more susceptible to changes in read length.

This study uses ONT data, paired with short-read data, as the ground truth, which better mitigates the impact of cellular heterogeneity compared to using bulk data. It also thoroughly explores the impact of different sequencing conditions on site identification and to demonstrate, from both gene and site perspectives, that differences in alignment and quantification tools can affect the conclusions of downstream analyses. This provides guidance for biologists in selecting appropriate methods for their research and offers insights for computational biologists in developing new methods.

Furthermore, although we simplified the results by using pseudobulk-level comparisons, this benchmark pipeline can be easily extended to different cell types (or clusters) and can quickly accommodate new APA analysis methods.

## 5 Methods

### 5.1 Data collection and preprocessing

Public data of single-cell RNA-seq (10x Chromium), spatial transcriptomics (10x Visium) data, and ONT data were collected from NCBI. The details of the data used in this paper can be found in **Table 1**. MOB C1 and E18 scRNA-seq data were based on the 10X Chromium Single Cell 3’ v2 reagent kit and MBM1 was based on the 10x Chromium Single Cell 3’ v3 reagent kit and all of them were quantified by the old version of Cell Ranger (Lebrigand, Magnone, et al., 2020; Tepe et al., 2018; Yang et al., 2022), which does not consider the counts from the intron. In order to facilitate the comparison of expression levels between genes and APAs which include the intronic forms, these data were re-analyzed by Cell Ranger (version 7.0.1).

E18, MOBV1, CBS1, and CBS2 consist of matched short-read and long-read data, meaning they share the same barcodes and UMIs (Lebrigand, Bergenstråhle, et al., 2023; Lebrigand, Magnone, et al., 2020). The original ONT data underwent processing using SiCeLoRe (Single Cell Long Read) version 2.1, resulting in a transcript-level expression matrix.

### 5.2 APA analysis tools

#### Identify polyA sites

As shown in **Figure 1A** and **Table S1**, the first category contains the most tools, involving different principles. We selected tools (scAPA, Sierra, SCAPTURE, SCAPE, MAAPER, scAPAtrap, polypipe) from each subcategory for further investigation. Following the respective tutorials for each tool, we configured the environment and executed these APA analysis tools to identify polyA sites across different datasets.

#### Annotate polyA sites

To minimize errors due to different annotation methods, we used the annotation function, AnnotationSite, provided by SCAPE to annotate the results produced by different methods. This function requires only the chromosome, site coordinate, and strand direction and can annotate sites up to 2kb downstream of the annotated 3’ UTR, making it applicable to all these tools. For methods that directly provide sites, those sites are used for annotation. For methods that provide peak intervals, if the interval is on the positive strand, the end position is retained; if the interval is on the negative strand, the start position is retained. After converting intervals to sites, further annotation was performed. To maintain consistency in annotation, the reference used for site annotation is derived from the Cell Ranger reference.

#### Evaluate resource usage

Runtime and maximum memory usage are widely used metrics for evaluating computational tools. We employed a Skylake CPU and tested three methods on MOB C1 and Mouse Brain E18 datasets, varying the number of cores from 2 to 32.

### 5.3 Comparison of identification performance

#### Evaluate site identification performance

To compare the overlap between sites, we first converted the ONT transcript matrix into a site matrix by extracting the chromosome, polyA site position, and strand direction. For transcripts on the positive strand, the polyA site corresponds to the transcript end position, while for transcripts on the negative strand, it corresponds to the transcript start position. Next, to ensure fairness, we only retained the polyA sites corresponding to genes identified by all APA analysis tools and ONT. Subsequently, we identified the number of overlapping sites in each gene using the findOverlap function in the R package GenomicRanges (version 1.46.1). We set the window size from 0 to 300nt with a step size of 10nt. Using strategy 2 (**Figure S2**), we counted the number of true positive sites based on the number of ground truth sites. The total number of true positive sites for each gene was then summed up to calculate precision and recall. To enhance the credibility of ONT data as ground truth and mitigate the impact of sequencing depth and quality, we calculated the coefficient of variation (CV) for each site. The CV was obtained by dividing the standard deviation of the expression level of each polyA site by its mean expression level. We set CV thresholds ranging from 2 to 32, with an increment of 2. This means that each CV threshold represents the retention of sites with a CV less than the threshold for performance comparison. We then repeated the aforementioned site identification performance evaluation steps for the retained sites to obtain precision and recall.

#### Downsampling analysis

To further identify the factors influencing site identification performance, we selected three representative methods (Sierra, scAPAtrap, and SCAPE) and applied them to the MOBC1 dataset to analyze whether changes in sequencing depth affect site identification. We used samtools (version 1.13.0, parameter -s, the value from 0.1 to 0.9) to downsample the BAM files from MOBC1 dataset. The MOBC1 dataset is derived from single-cell sequencing of the mouse olfactory bulb, containing five million reads, suitable for downstream analysis through subsampling. The generated downsampled BAM files contain 10%, 30%, 50%, 70%, or 90% of all reads and were used as input for APA analysis tools to reidentify sites. We then compared how the number of sites changed for different methods as the coverage decreased.

#### Read length truncation analysis

Similarly, we applied these three methods to the MBM1 dataset to examine whether read length affects site identification. MBM1 dataset is the scRNA-seq of the mouse brain with read lengths reaching 150 nucleotides. We truncated its reads using Cell Ranger (version 7.0.1, parameter –r2-length, from 120 to 30) and remapped and quantified them. BAM files from mappings with different lengths were used for the re-identification of polyA sites, and differences in site identification among different methods were compared as the length was shortened.

#### Compare splicing awareness among methods by the coverage plot

Among the three methods, scAPAtrap is not splice-aware and Sierra is splice-aware, but it remains uncertain whether SCAPE can effectively differentiate false positive peaks caused by splicing. Utilizing coverage plots provides an intuitive and effective means to detect false positive peaks resulting from splicing events. By examining coverage variations at the single-base level and comparing the disparities in identified positions across different methods, we assess the extent to which splicing influences site identification. Coverage plots were drawn by Gviz package (version 1.42.1).

#### Distribution of identified sites and its corresponding genes

We conducted a statistical analysis of the distribution of polyA sites. First, we calculated the number of genes corresponding to different numbers of sites. To mitigate the potential scaling effect of multi-site genes, we categorized genes with more than six loci into a single group and visualized the distribution using a histogram generated by ggplot2. Next, based on the annotation results, we analyzed the distribution of different annotation categories (exon, intron, UTR, and downstream) under different site counts. For example, we determined how many sites identified in genes with two sites originated from exons, introns, or UTR regions. Finally, we identified the overlap in genes identified by each method and visualized the gene overlap using an upset plot using the R package UpSetR (version 1.4.0).

### 5.4 Comparison of quantification performance

#### Evaluate site quantification performance

Similar to the site identification steps, the quantification of site expression begins with identifying overlapping sites for all genes. Subsequently, the findOverlap function is used to identify overlapping sites within a given window range. The read count for each site is normalized by the total number of reads within the corresponding cell (and times 10,000 to obtain normalized values). The Pearson correlation coefficient and relative error of the overlapping site are calculated to compare the quantification performance. The relative error is defined as the absolute error of expression level between ONT sites and the predicted sites divided by the ONT expression level. To eliminate the impact of identification discrepancies, we further filtered sites that overlapped across all methods and ONT, and then recalculated the correlation coefficient and relative error to analyze whether differences in quantification steps among these methods would result in significant differences in quantification outcomes.

#### Evaluate gene quantification performance

We merged sites from the same gene and sum up UMI counts. Raw counts were divided by the total counts and times 10,000 to obtain normalized values. Similarly, we selected three representative methods to analyze whether the quantification performance at the gene level would affect the quantification at the site level. This analysis is divided into two parts. First, we compared the correlation of gene expression between different methods, particularly the correlation between ONT gene expression and the quantification results from Cell Ranger or Space Ranger. This comparison demonstrates the validity of using ONT as the ground truth for quantification at the site level. Second, we compared the expression levels obtained from different APA analysis methods with the quantification results from Cell Ranger or Space Ranger. Theoretically, APA analysis tools use BAM files generated by Cell Ranger or Space Ranger as input for identification and quantification. Therefore, the gene expression levels aggregated from sites identified by these tools should not only exhibit good correlation but also should not exceed the gene expression levels obtained by Ranger tools. Consequently, for each gene and each cell, we identified whether the quantification results of each method were consistent with their corresponding Ranger quantification results and whether there were any outliers, i.e., cases where the expression levels exceeded those of Ranger tools. These results were visualized using scatter plots generated by ggplot2 (version 3.5.1).

### 5.5 Comparison of differential expression performance

#### Evaluation of Differentially Expressed Site Identification Performance

We selected the two cell types with the highest number of cells and identified differentially expressed sites between these two cell types. To enhance the effectiveness of differential expression identification, we employed a pseudobulk strategy, randomly dividing each cell group into three replicates, and then used DESeq2 (version 1.34.0) to identify differentially expressed sites. Sites with an FDR less than 0.05 were retained. Subsequently, we compared the differentially expressed sites predicted by the tools with those identified by ONT. Similar to the identification process, we used different findOverlap functions to calculate the number of overlapping sites. Since the sequencing depth of ONT is generally lower than that of corresponding Illumina sequencing, the number of sites obtained by ONT is significantly fewer than those obtained by APA analysis tools. Therefore, when selecting metrics, we chose to calculate recall and the correlation coefficient of logFC to evaluate the performance of differential expression analysis.

#### Evaluate differentially expressed genes

We also analyzed the differential gene expression between the two cell types. Using DESeq2 integrated within Seurat, we identified differentially expressed genes and calculated recall and the correlation coefficient of logFC.

## 6 Declarations

### 6.1 Ethics approval and consent to participate

Not applicable

### 6.2 Consent for publication

Not applicable

### 6.3 Availability of data and materials

The publicly available data used for analysis are available in the following GEO: GSE130708, GSE153859, GSE121891, GSE169606. Benchmark code for this research is available in the https://github.com/qianzhao613/APA_benchmark_tests.

### 6.4 Competing interests

The authors declare no competing interests.

### 6.5 Funding

MR was supported by a Wellcome Trust Discovery Award (Ref 227415/Z/23/Z). QZ was supported by a University of Manchester-China Scholarship Council Joint Scholarship.

### 6.6 Authors’ contributions

MR and QZ conceptualized the study. QZ performed data analysis and wrote the manuscript. MR reviewed the manuscript.

## 6.7 Acknowledgements

We thank the authors of the datasets used in this study. We also appreciate the constructive discussions with all members of the Rattray Lab.

## Supplementary Information

### Supplemental Figures

**Figure S1:**
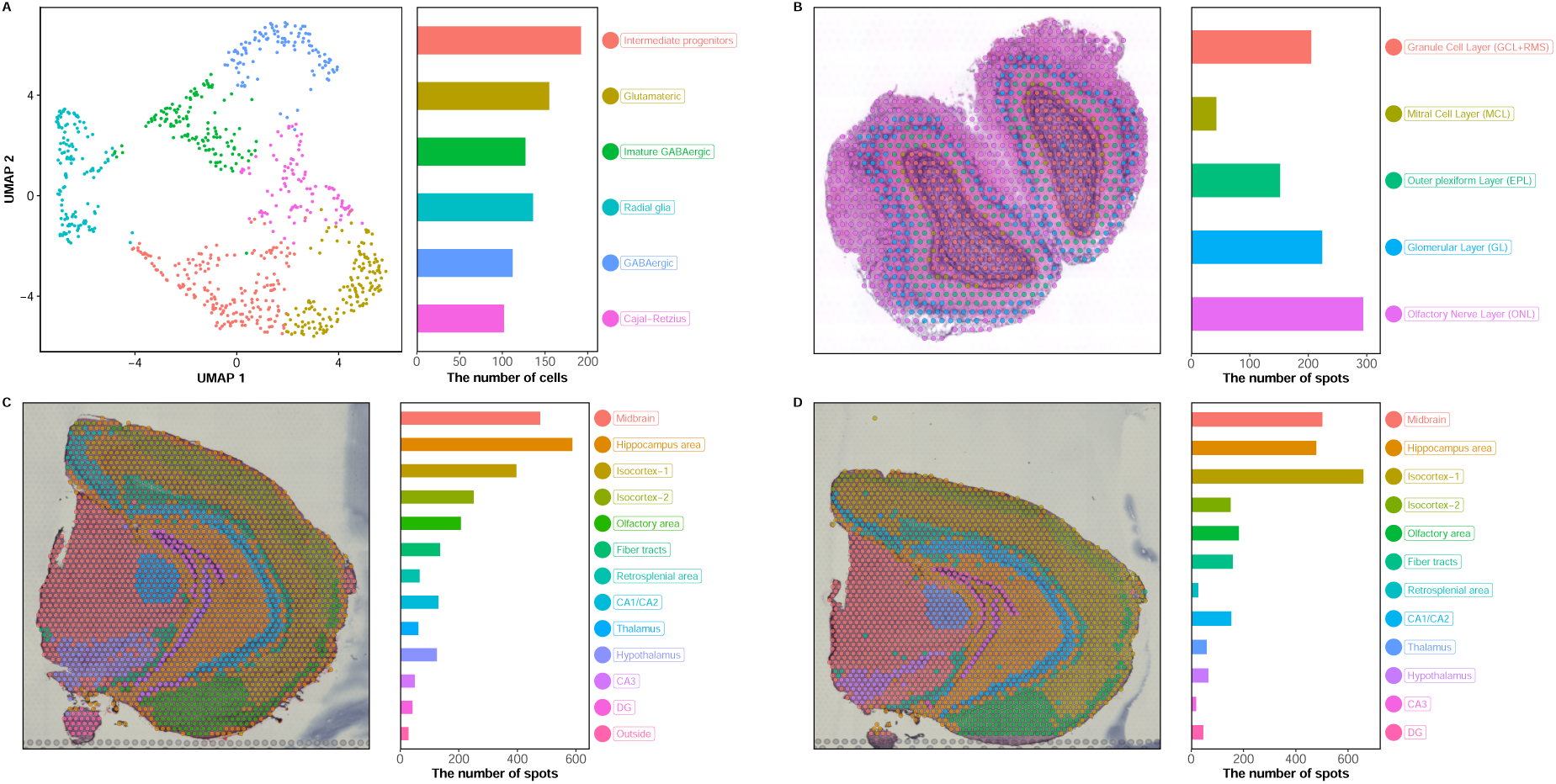
Cluster information of datasets. **(A)** Clustering results for the single-cell dataset E18. The left panel shows a UMAP plot, while the right panel presents a bar chart depicting the number of cells contained within each cell type, with different colours representing different cell types. **(B)** Clustering results for the spatial transcriptomics dataset MOBV1. The left panel displays the spatial locations on the HE image, while the right panel features a bar chart illustrating the number of spots corresponding to different regions, with different colours representing different regions. **(C)** Clustering results for the spatial transcriptomics dataset CBS1. **(D)** Clustering results for the spatial transcriptomics dataset CBS2.

**Figure S2:**
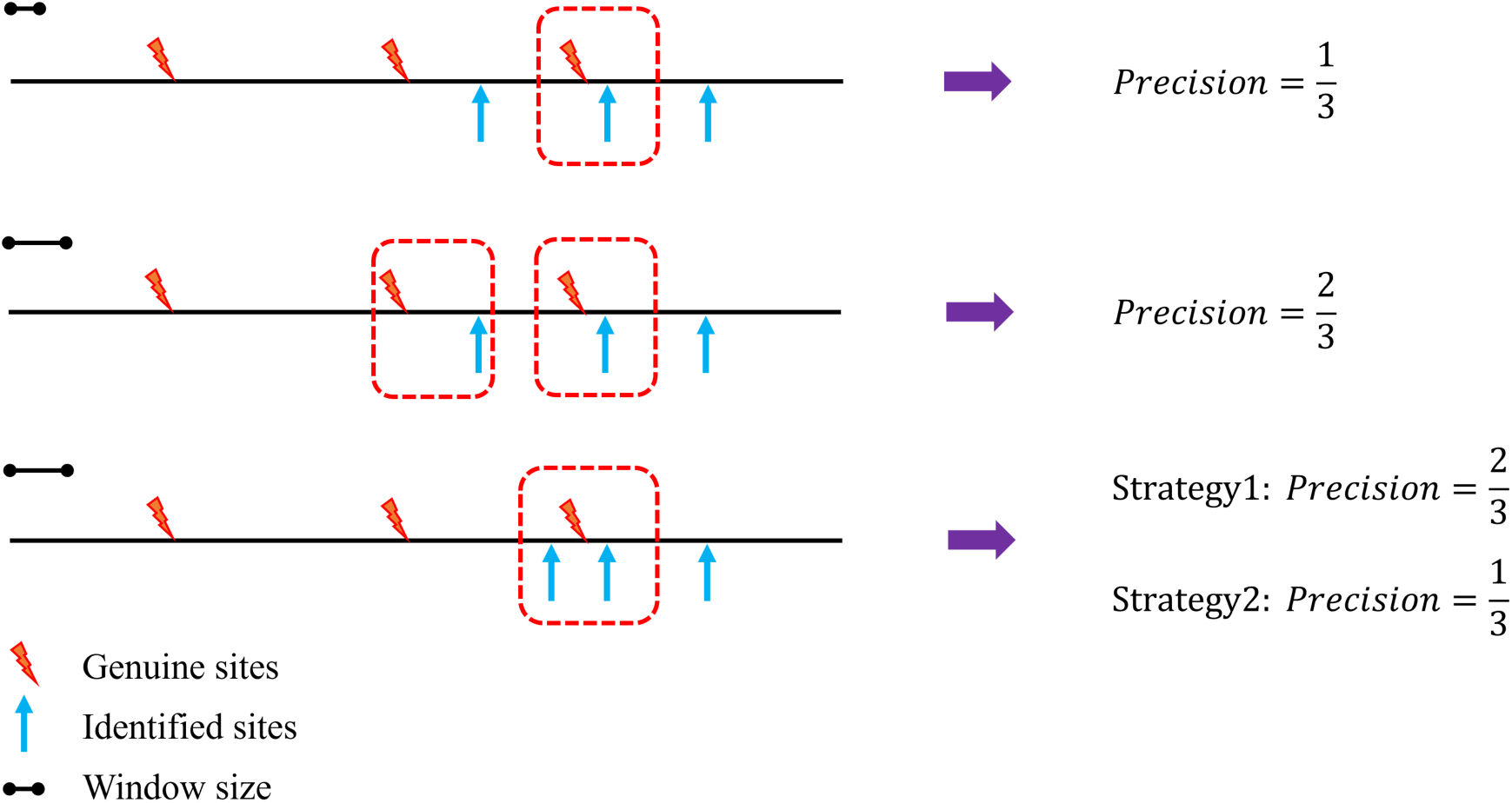
Strategies for calculating the number of true positive sites. Schematic showing that the hypothetical gene contains three genuine polyA sites and that the APA analysis tool identifies three sites. The top row shows that when the window is small, a single predicted site falls within the window interval of the third true site, and other sites do not overlap. Therefore, the true positive site is 1, and the precision is one-third. The window in the middle row is expanded so that the first identified site also coincides with the second true site. The number of true positive sites becomes 2, and the precision is equal to two-thirds. The bottom row shows the special case where two identified sites fall within the window interval of the same ground-truth site. In this case, two strategies arise for calculating the number of true positives. In the first strategy, both identified sites are counted as coinciding with the true sites, so the number of true positive sites is 2 and the precision is two-thirds. The second strategy believes that it should be calculated based on the number of ground-truth sites, so the true positive site is 1 and the precision is one-third.

**Figure S3:**
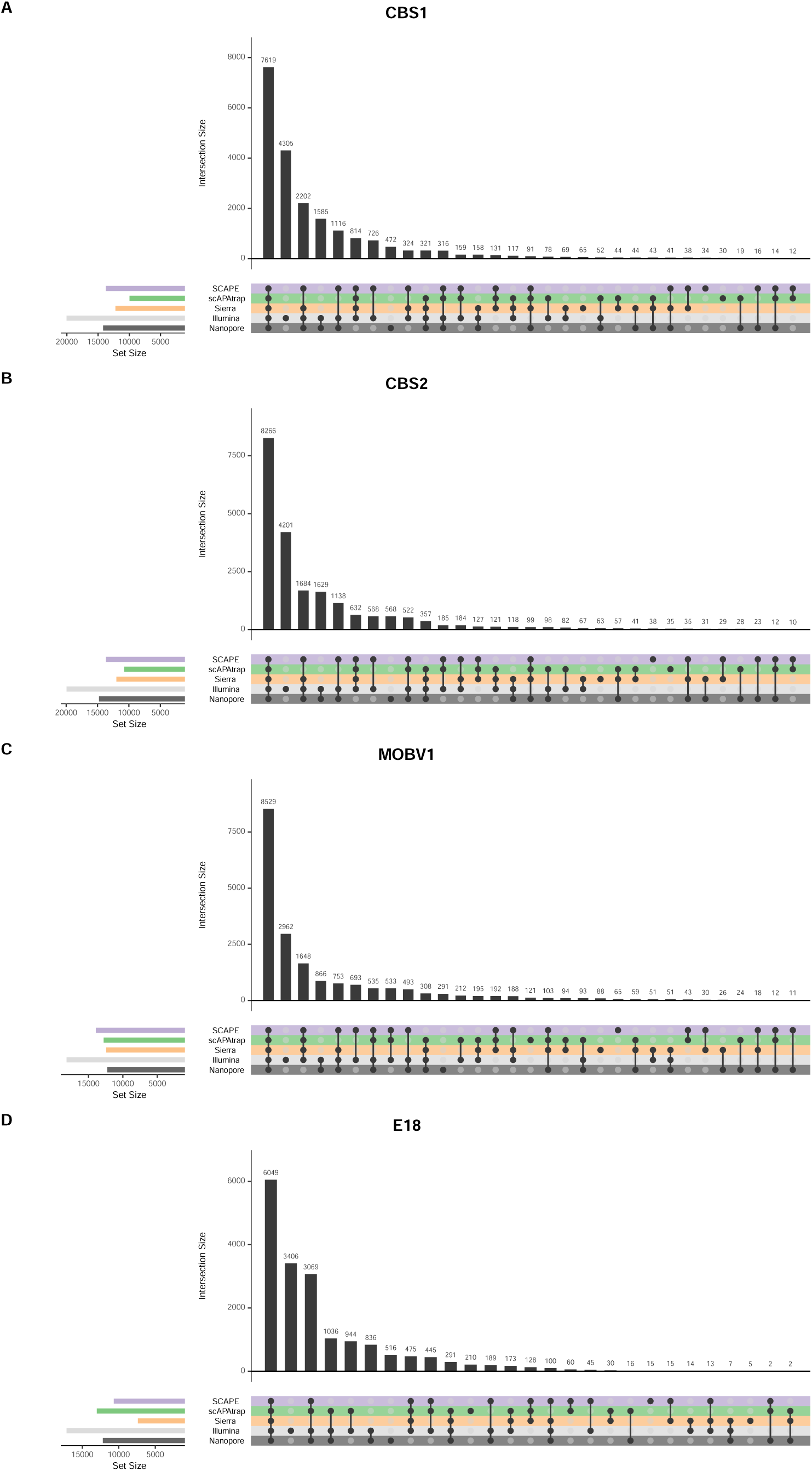
Overlapped genes between different methods among four datasets. **(A)** CBS1. **(B)** CBS2. **(C)** MOBV1 **(D)** E18

**Figure S4:**
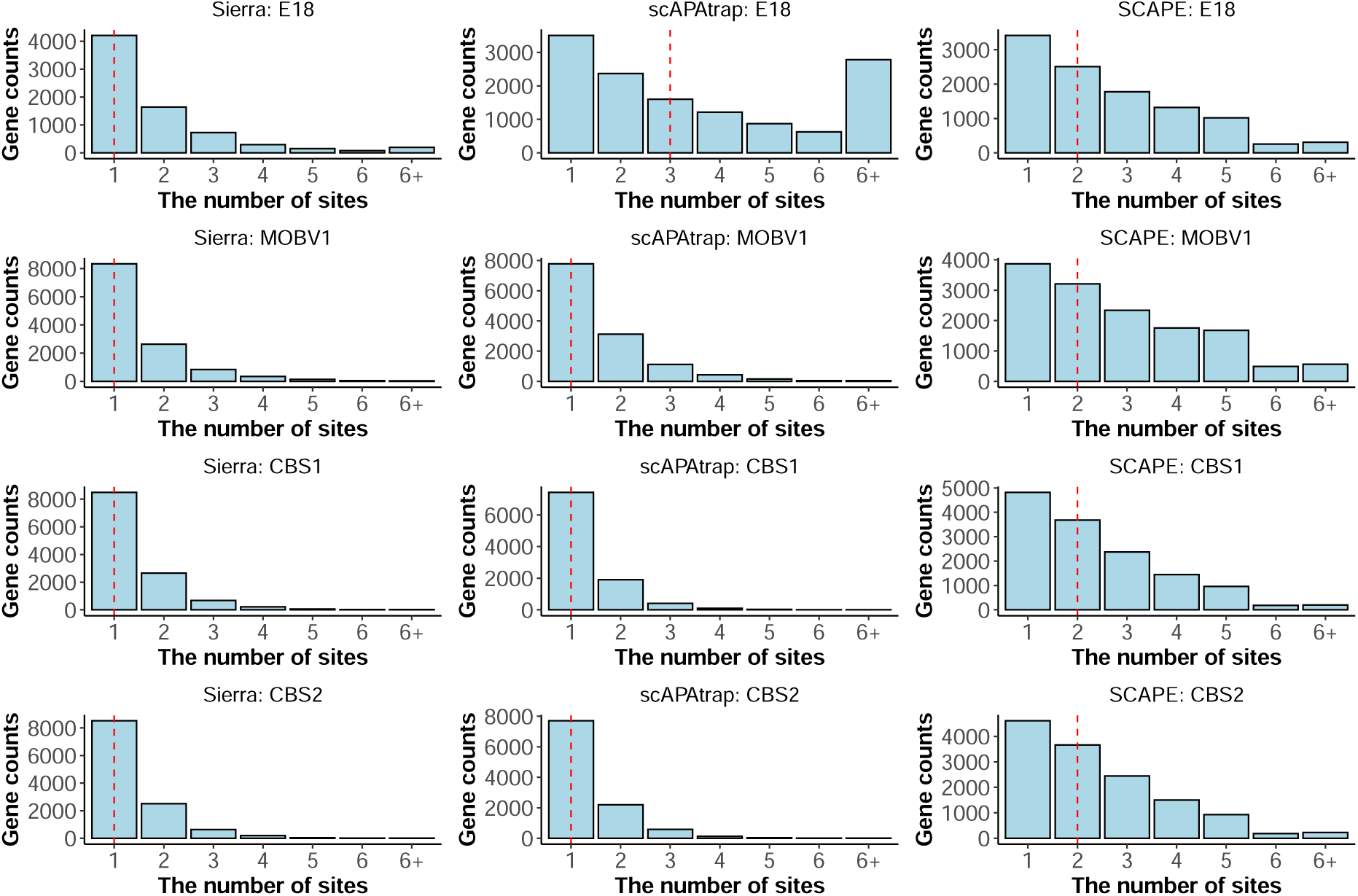
The distribution of the number of sites. Categorization is based on the number of identified sites per gene. The x-axis represents the number of sites associated with each gene, while the y-axis indicates the number of genes. The red dashed line represents the median. Genes with more than six sites are considered as a distinct category.

**Figure S5:**
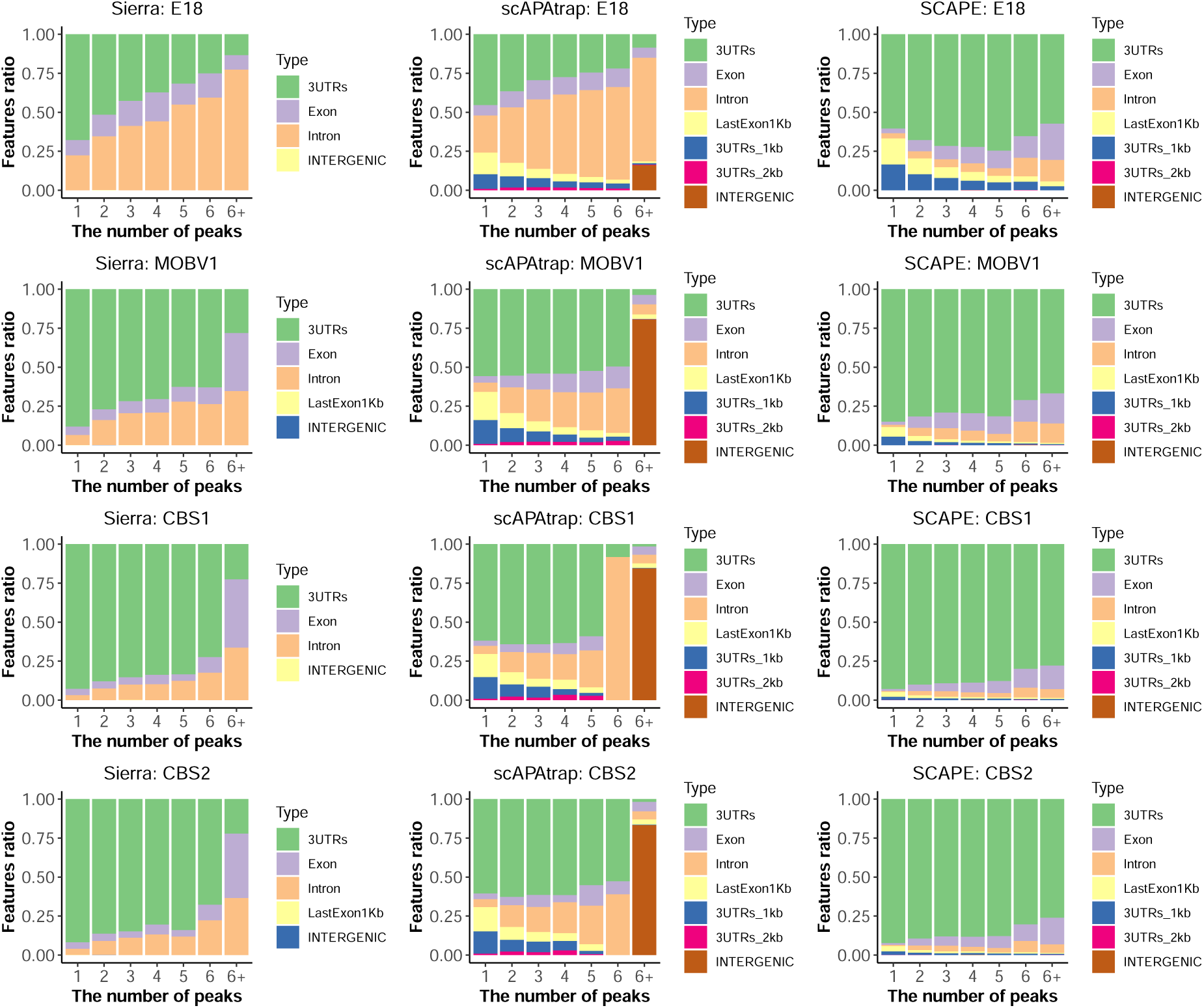
The distribution of genomic features. Annotated sites were classified into seven categories based on their overlap with genomic features: 3’ UTR, Exon, Intron, Intergenic, Last Exon 1kb, 3’ UTR 1kb, and 3’ UTR 2kb. Different colors represent different genomic features.

**Figure S6:**
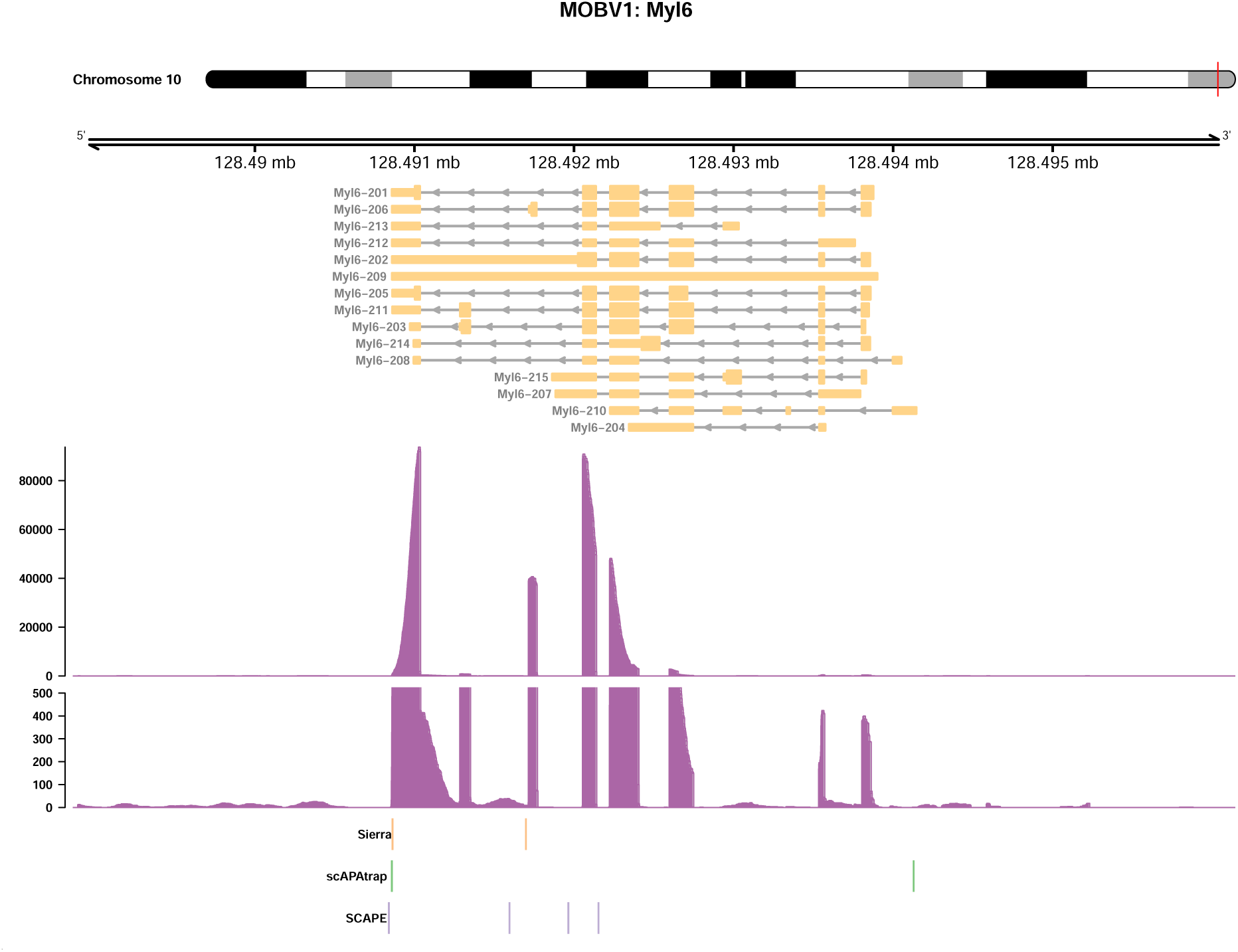
Splicing influence identification. Coverage of the Myl6 gene in the MOBV1 dataset. At the top is the corresponding chromosomal location, followed by transcripts from the reference annotation, the per-base read coverage (including subplot ranging from 0 to 500, to illustrate potential low-expression isoforms), and the locations of the sites identified by the three APA analysis methods.

**Figure S7:**
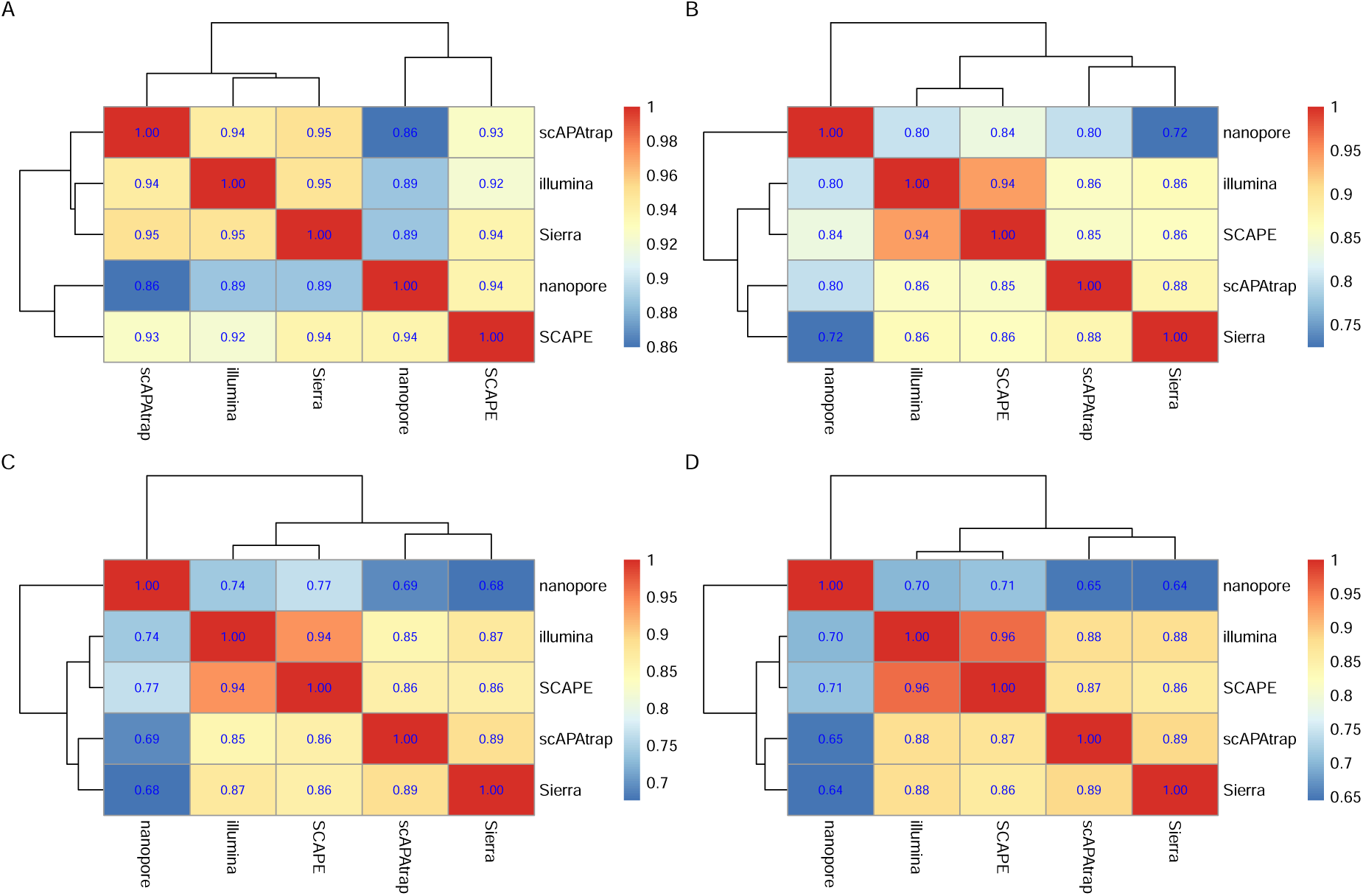
Gene expression correlation heatmap. **(A)** E18 Correlation between gene expression levels obtained by APA analysis methods and those obtained by Cell Ranger, Space Ranger, and ONT. **(B)** CBS1 Correlation between gene expression levels obtained by APA analysis methods and those obtained by Cell Ranger, Space Ranger, and ONT. **(C)** CBS2 Correlation between gene expression levels obtained by APA analysis methods and those obtained by Cell Ranger, Space Ranger, and ONT. **(D)** MOBV1 Correlation between gene expression levels obtained by APA analysis methods and those obtained by Cell Ranger, Space Ranger, and ONT.

**Figure S8:**
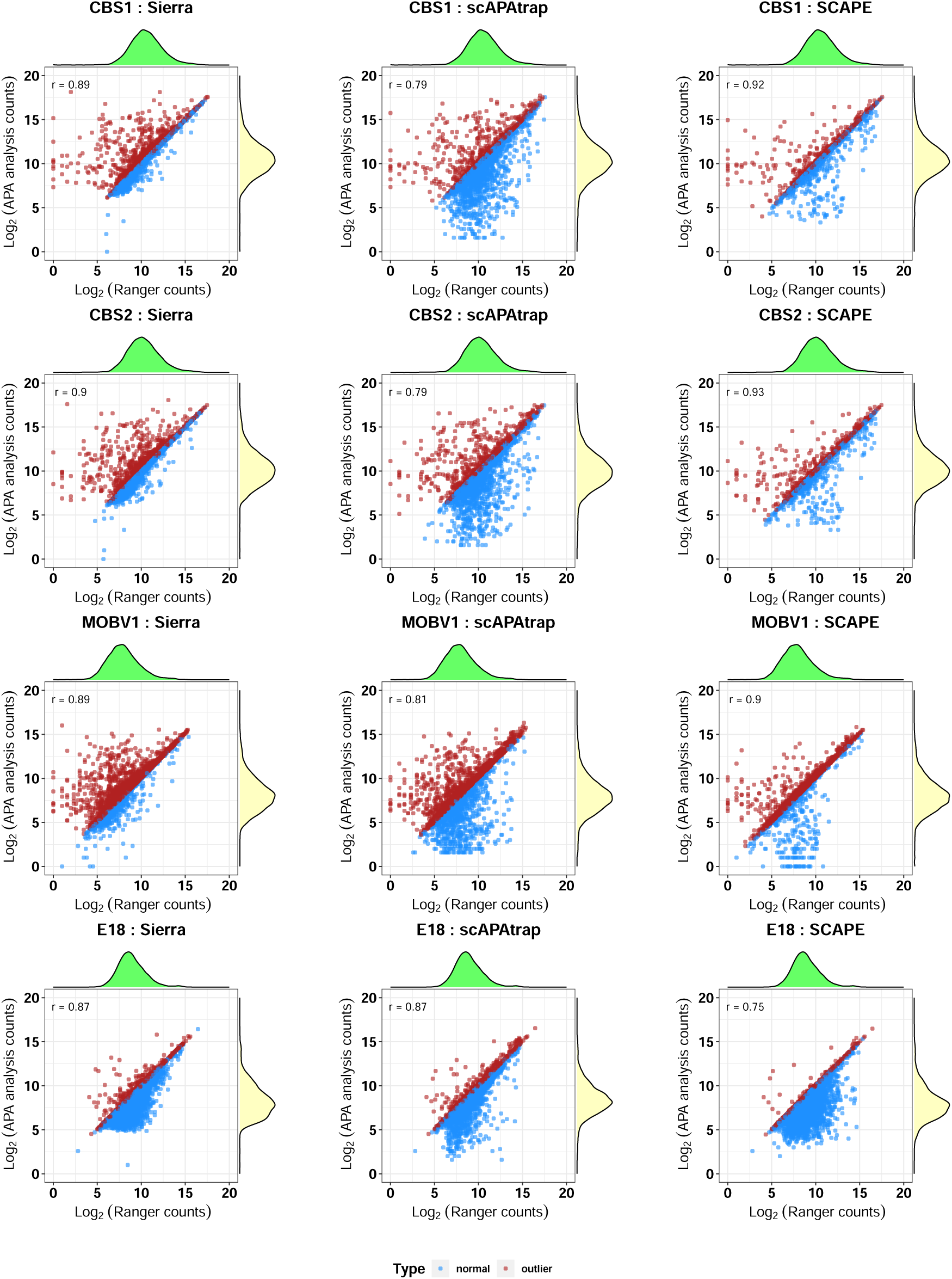
Gene expression scatter plot. Correlation between gene expression levels obtained by APA analysis methods and those obtained by Cell Ranger or Space Ranger. Each point represents a gene, with the x-axis showing the log-transformed gene expression levels from Cell Ranger or Space Ranger, and the y-axis showing the log-transformed expression levels obtained by the APA analysis tool. Red dots indicate outliers, where the APA tool’s expression levels exceed those of Cell Ranger or Space Ranger, while blue dots represent normal data.

**Figure S9:**
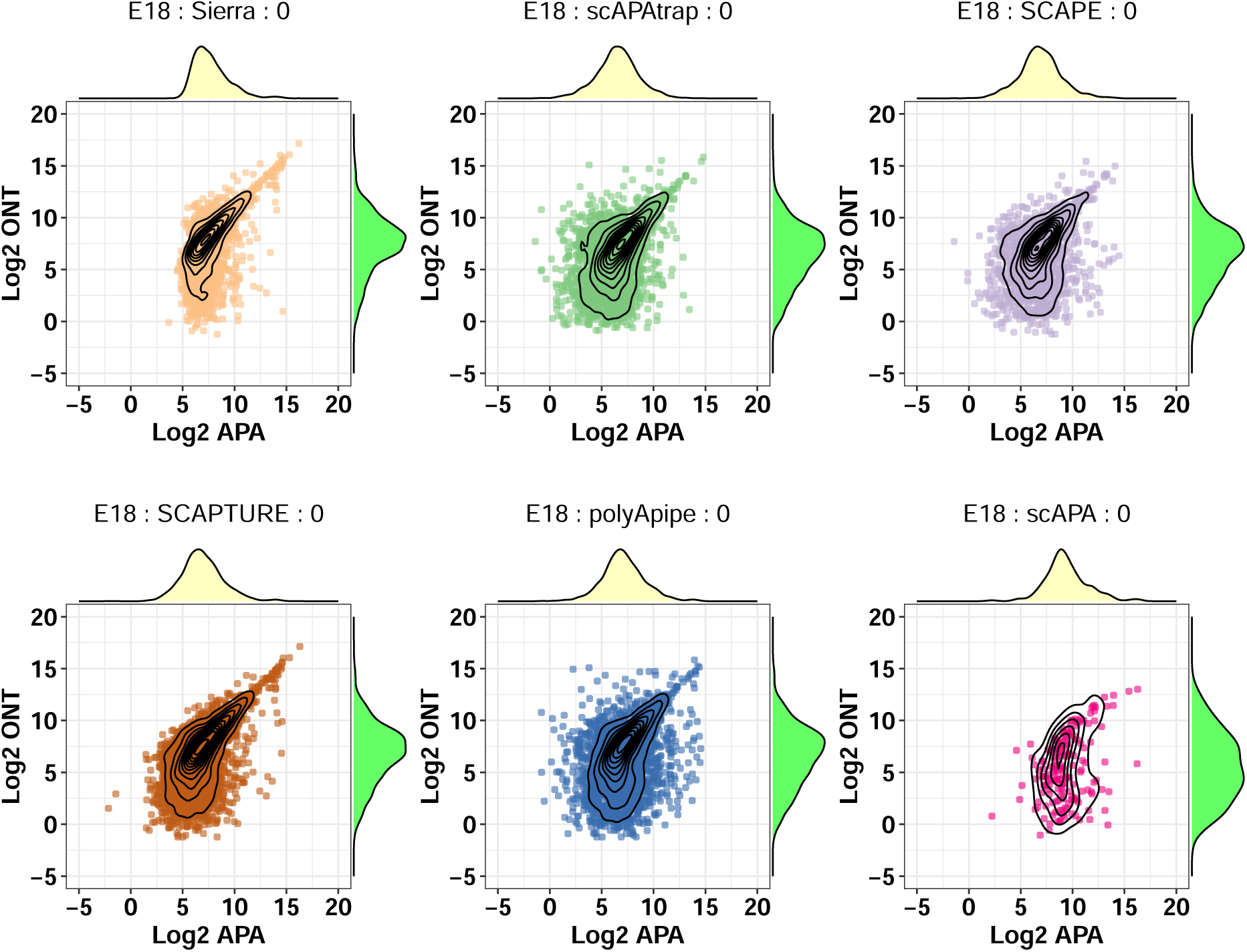
Site scatter plot at 0nt window size. Correlation between APA analysis methods and ONT at 0nt window size. Each point represents a gene, with the x-axis showing the log-transformed gene expression levels from Cell Ranger or Space Ranger, and the y-axis showing the log-transformed expression levels obtained by the APA analysis tool.

**Figure S10:**
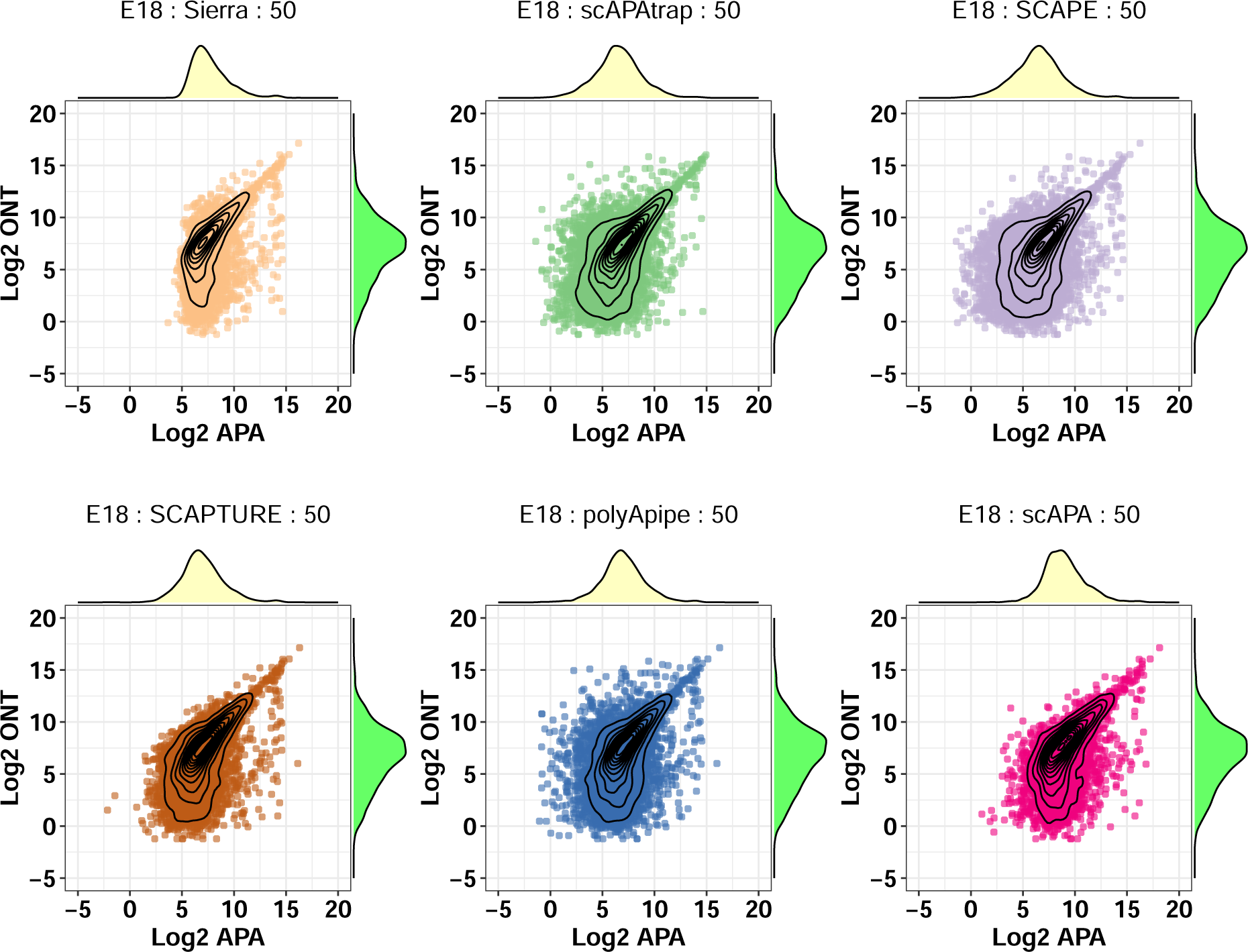
Site scatter plot at 50nt window size. Correlation between gene expression levels obtained by APA analysis methods and those obtained by Cell Ranger or Space Ranger. Each point represents a gene, with the x-axis showing the log-transformed gene expression levels from Cell Ranger or Space Ranger, and the y-axis showing the log-transformed expression levels obtained by the APA analysis tool.

**Figure S11:**
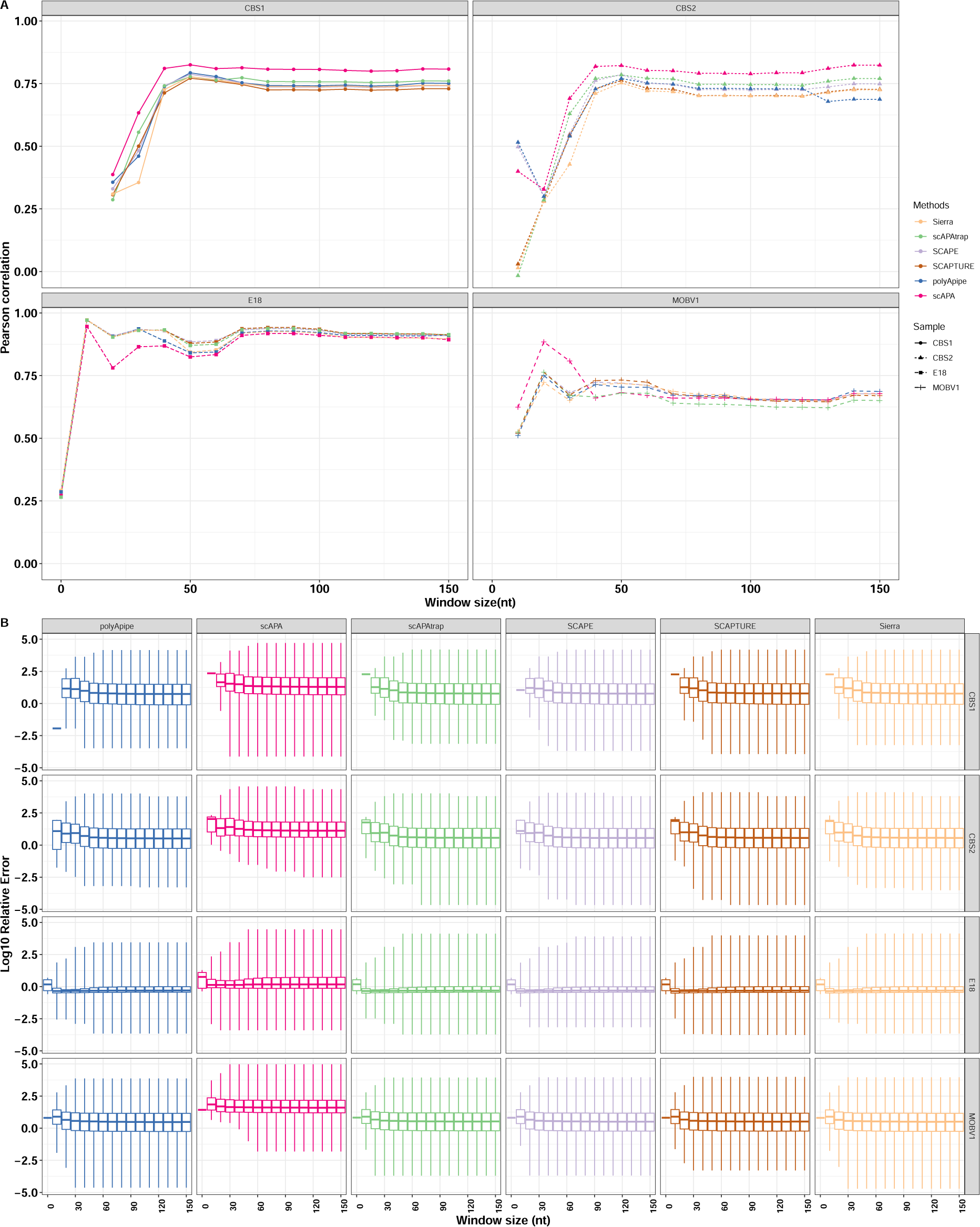
Quantification performance on the same set of sites. **(A)** Pearson correlation on the same set of sites. Common sites were obtained through the intersection of ONT sites. **(B)** RE on the same set of sites.

### Supplemental Tables

**Table S1:** Categories of APA analysis tools.

| Method | Category | Subcategory | Language | Version | Reference | GitHub |
| --- | --- | --- | --- | --- | --- | --- |
| scAPA | 1 | 1 | R | na | Shulman and Elkon, 2019 | https://github.com/ElkonLab/scAPA |
| Sierra | 1 | 1 | R | na | Patrick et al., 2020 | https://github.com/VCCRI/Sierra |
| SCAPTURE | 1 | 1 | Shell + Python | 1.1.1 | G.-W. Li et al., 2021 | https://github.com/YangLab/SCAPTURE |
| MAAPER | 1 | 2 | R | 1.1.1 | W. V. Li et al., 2021 | https://github.com/Vivianstats/MAAPER |
| SCAPE | 1 | 2 | Python+R | na | Zhou et al., 2022 | https://github.com/LuChenLab/SCAPE |
| scAPAttrap | 1 | 3 | R | 0.1.0 | Wu et al., 2021 | https://github.com/BMILAB/scAPAttrap |
| scDarPars | 1 | 3 | Python | 0.0.1 | Gao et al., 2021 | https://github.com/YiPeng-Gao/scDaPars |
| polyApipe | 1 | 4 | R+Python | 0.1.0 | Unpublished | https://github.com/MonashBioinformaticsPlatform/polyApipe |
| tail anchor from scAPAttrap | 1 | 4 | R | 0.1.0 | Wu et al., 2021 | https://github.com/BMILAB/scAPAttrap |
| SCINPAS | 1 | 4 | Python+Groovy (Nextflow) | 1.0.0 | Moon, Burri, and Zavolan, 2023 | https://github.com/zavolanlab/SCINPAS |

## Notes

### Competing Interest Statement

The authors have declared no competing interest.

## References

Bryce-Smith, Sam et al. (June 26, 2023). Extensible benchmarking of methods that identify and quantify polyadenylation sites from RNA-seq data. Pages: 2023.06.23.546284 Section: New Results. doi: 10.1101/2023.06.23.546284.

Fansler, Mervin M., Sibylle Mitschka, and Christine Mayr (June 22, 2023). Comprehensive annotation of 3’UTRs from primary cells and their quantification from scRNA-seq data. Pages: 2021.11.22.469635 Section: New Results. doi: 10.1101/2021.11.22.469635.

Gao, Yipeng, et al. (Oct. 2021). “Analysis of alternative polyadenylation from single-cell RNA-seq using scDaPars reveals cell subpopulations invisible to gene expression”. In: Genome Research 31.10, pp. 1856–1866. issn: 1549-5469. doi: 10.1101/gr.271346.120.

Gillen, Austin E., Raeann Goering, and J. Matthew Taliaferro (2021). “Quantifying alternative polyadenylation in RNAseq data with LABRAT”. In: Methods in Enzymology 655, pp. 245–263. issn: 1557-7988. doi: 10.1016/bs.mie.2021.03.018.

Goering, Raeann et al. (June 26, 2021). “LABRAT reveals association of alternative polyadenylation with transcript localization, RNA binding protein expression, transcription speed, and cancer survival”. In: BMC Genomics 22.1, p. 476. issn: 1471-2164. doi: 10.1186/s12864-021-07781-1.

Kowalski, Madeline H. et al. (Feb. 10, 2023). CPA-Perturb-seq: Multiplexed single-cell characterization of alternative polyadenylation regulators. Pages: 2023.02.09.527751 Section: New Results. doi: 10.1101/2023.02.09.527751.

Lebrigand, Kevin, Joseph Bergenstråhle, et al. (May 8, 2023). “The spatial landscape of gene expression isoforms in tissue sections”. In: Nucleic Acids Research 51.8, e47. issn: 1362-4962. doi: 10.1093/nar/gkad169.

Lebrigand, Kevin, Virginie Magnone, et al. (Aug. 12, 2020). “High throughput error corrected Nanopore single cell transcriptome sequencing”. In: Nature Communications 11, p. 4025. issn: 2041-1723. doi: 10.1038/s41467-020-17800-6.

Li, Guo-Wei et al. (Aug. 10, 2021). “SCAPTURE: a deep learning-embedded pipeline that captures polyadenylation information from 3’ tag-based RNA-seq of single cells”. In: Genome Biology 22.1, p. 221. issn: 1474-760X. doi: 10.1186/s13059-021-02437-5.

Li, Wei Vivian et al. (Aug. 10, 2021). “MAAPER: model-based analysis of alternative polyadenylation using 3’ end-linked reads”. In: Genome Biology 22.1, p. 222. issn: 1474-760X. doi: 10.1186/s13059-021-02429-5.

Meyer, Elisabeth et al. (Oct. 25, 2022). “ReadZS detects cell type-specific and developmentally regulated RNA processing programs in single-cell RNA-seq”. In: Genome Biology 23.1, p. 226. issn: 1474-760X. doi: 10.1186/s13059-022-02795-8.

Moon, Youngbin, Dominik Burri, and Mihaela Zavolan (Sept. 2023). “Identification of experimentally-supported poly(A) sites in single-cell RNA-seq data with SCINPAS”. In: NAR Genomics and Bioinformatics 5.3, lqad079. issn: 2631-9268. doi: 10.1093/nargab/lqad079. eprint: https://academic.oup.com/nargab/article-pdf/5/3/lqad079/51517239/lqad079.pdf.

Pan, Lu et al. (Mar. 1, 2022). “Isoform-level quantification for single-cell RNA sequencing”. In: Bioinformatics 38.5, pp. 1287–1294. issn: 1367-4803. doi: 10.1093/bioinformatics/ btab807.

Patrick, Ralph et al. (July 8, 2020). “Sierra: discovery of differential transcript usage from polyA-captured single-cell RNA-seq data”. In: Genome Biology 21.1, p. 167. issn: 1474-760X. doi: 10.1186/s13059-020-02071-7.

Shulman, Eldad David and Ran Elkon (Nov. 4, 2019). “Cell-type-specific analysis of alternative polyadenylation using single-cell transcriptomics data”. In: Nucleic Acids Research 47.19, pp. 10027–10039. issn: 1362-4962. doi: 10.1093/nar/gkz781.

Tepe, Burak et al. (Dec. 4, 2018). “Single-Cell RNA-Seq of Mouse Olfactory Bulb Reveals Cellular Heterogeneity and Activity-Dependent Molecular Census of Adult-Born Neurons”. In: Cell reports 25.10, 2689–2703.e3. issn: 2211-1247. doi: 10.1016/j.celrep.2018.11. 034.

Wu, Xiaohui et al. (July 20, 2021). “scAPAtrap: identification and quantification of alternative polyadenylation sites from single-cell RNA-seq data”. In: Briefings in Bioinformatics 22.4, bbaa273. issn: 1477-4054. doi: 10.1093/bib/bbaa273.

Yang, Chunying et al. (May 11, 2022). “Single-cell transcriptomics identifies premature aging features of TERC-deficient mouse brain and bone marrow”. In: GeroScience 44.4, pp. 2139–2155. issn: 2509-2715. doi: 10.1007/s11357-022-00578-4.

Ye, Wenbin, et al. (Feb. 2023). “A Survey on Methods for Predicting Polyadenylation Sites from DNA Sequences, Bulk RNA-seq, and Single-cell RNA-seq”. In: Genomics, Proteomics & Bioinformatics 21.1, pp. 67–83. issn: 16720229. doi: 10.1016/j.gpb.2022.09.005.

Zhou, Ran et al. (June 24, 2022). “SCAPE: a mixture model revealing single-cell polyadenylation diversity and cellular dynamics during cell differentiation and reprogramming”. In: Nucleic Acids Research 50.11, e66. issn: 1362-4962. doi: 10.1093/nar/gkac167.

